# Reversing age: dual species measurement of epigenetic age with a single clock

**DOI:** 10.1101/2020.05.07.082917

**Authors:** Steve Horvath, Kavita Singh, Ken Raj, Shraddha Khairnar, Akshay Sanghavi, Agnivesh Shrivastava, Joseph A. Zoller, Caesar Z. Li, Claudia B. Herenu, Martina Canatelli-Mallat, Marianne Lehmann, Leah C. Solberg Woods, Angel Garcia Martinez, Tengfei Wang, Priscila Chiavellini, Andrew J. Levine, Hao Chen, Rodolfo G. Goya, Harold L. Katcher

**Affiliations:** Department of Human Genetics, David Geffen School of Medicine, University of California, Los Angeles, Los Angeles, California, USA; Department of Biostatistics, Fielding School of Public Health, University of California, Los Angeles, Los Angeles, California, USA; Shobhaben Pratapbhai Patel School of Pharmacy and Technology Management, SVKM’S NMIMS University, Mumbai, India; Radiation Effects Department, Centre for Radiation, Chemical and Environmental Hazards, Public Health England, Chilton, Didcot, UK; Nugenics Research Pvt Ltd, India; Institute for Experimental Pharmacology of Cordoba (IFEC), School of Chemical Sciences, National University of Cordoba, Cordoba, Argentina; Biochemistry Research Institute of La Plata – Histology B, Pathology B, School of Medicine, University of La Plata, La Plata CC 455 (zip 1900), Argentina; Wake Forest University School of Medicine, 1 Medical Center Drive, Winston Salem, NC 27157, USA; Department of Pharmacology, Addiction Science and Toxicology, The University of Tennessee Health Science Center, Memphis, TN 3993, USA; Department of Neurology, David Geffen School of Medicine at the University of California, Los Angeles, CA, 90095, USA

**Keywords:** rejuvenation, plasma fraction, epigenetic clock, DNA methylation, rat

## Abstract

Young blood plasma is known to confer beneficial effects on various organs in mice. However, it was not known whether young plasma rejuvenates cells and tissues at the epigenetic level; whether it alters the epigenetic clock, which is a highly-accurate molecular biomarker of aging. To address this question, we developed and validated six different epigenetic clocks for rat tissues that are based on DNA methylation values derived from n=593 tissue samples. As indicated by their respective names, the rat pan-tissue clock can be applied to DNA methylation profiles from all rat tissues, while the rat brain-, liver-, and blood clocks apply to the corresponding tissue types. We also developed two epigenetic clocks that apply to both human and rat tissues by adding n=850 human tissue samples to the training data. We employed these six clocks to investigate the rejuvenation effects of a plasma fraction treatment in different rat tissues. The treatment more than halved the epigenetic ages of blood, heart, and liver tissue. A less pronounced, but statistically significant, rejuvenation effect could be observed in the hypothalamus. The treatment was accompanied by progressive improvement in the function of these organs as ascertained through numerous biochemical/physiological biomarkers and behavioral responses to assess cognitive functions. Cellular senescence, which is not associated with epigenetic aging, was also considerably reduced in vital organs. Overall, this study demonstrates that a plasma-derived treatment markedly reverses aging according to epigenetic clocks and benchmark biomarkers of aging.

## INTRODUCTION

Several decades ago, heterochronic parabiosis studies, in which the circulations of old and young mice were connected to instigate mixing of their blood, revealed that young blood can rejuvenate aging mice and vice versa – old blood ages young mice ^1^. A recent revival of this approach confirmed this observation and demonstrated beneficial effects on muscle, heart, brain, and numerous other organs ^2–7^. Repeated injections of young plasma into older mice, an effective alternative to parabiosis, confirmed the prevailing notion that the beneficial effect is due to blood-borne factors; as opposed to the possibility that old mice might have, by proxy benefited from better-functioning organs of the young ^8^. Many investigations and commercial investments have been made to identify and isolate from blood, the rejuvenation factor/s, which can in theory, be of benefit to mitigate or treat age-related conditions such as Alzheimer’s disease and more. The plasma fraction treatment, used in the investigation described below, is based on the principle of Heterochronic Plasma Exchange (HPE), whereby the plasma of old rats is replaced by those from the young ^9^. This approach is derived from heterochronic parabiosis, but without the need to physically attach the circulatory systems of two animals together. In addition to greatly reducing the stress to the animals, HPE is expected to have a more profound effect as 100% of the old animal’s blood could be replaced. This is in contrast to heterochronic parabiosis, where a young rat, with approximately half the weight of an old rat, contributes less than 50% to the combined plasma circulation in the parabiotic partners. The general loss of tissue repair with age may reflect the negative influence of age-accumulated inhibitory proteins in aged tissues and circulation ^10^. Therefore, if putative pro-aging factors are present in the blood plasma of old animals ^10^, they would remain in the parabiotic partner.

When considering the concept of aging and rejuvenation, it is important to appreciate that improved health or organ function through medication or surgery does not necessarily indicate molecular age reversal. Hence, it is conceptually challenging to test whether plasma fraction treatment, or any other putative treatment, actually reverses biological age, because there is no consensus on how to measure biological aging ^11^. We addressed this challenge by using both clinical biomarkers and molecular biomarkers of aging. While clinical biomarkers have obvious advantages (being indicative of organ dysfunction or disease), they are neither sufficiently mechanistic nor proximal to fundamental mechanisms of aging to serve as indicators of them. It has long been recognized that epigenetic changes are one of several primary hallmarks of aging ^12–16^. With the technical advancement of methylation array platforms that can provide quantitative and accurate profiles of specific CpG methylations; came the insight to combine methylation levels of several DNA loci to develop an accurate age estimator ^17–21^. Such DNA methylation (DNAm) age estimators exhibit unexpected properties: they apply to all sources of DNA (sorted cells, tissues, and organs) and surprisingly to the entire age spectrum (from prenatal tissue samples to tissues of centenarians) ^20,22^. A substantial body of literature demonstrates that these epigenetic clocks capture aspects of biological age ^22^. This is demonstrated by the finding that the discrepancy between DNAm age and chronological age (term as “epigenetic age acceleration”) is predictive of all-cause mortality even after adjusting for a variety of known risk factors ^23–25^. Pathologies and conditions that are associated with epigenetic age acceleration includes, but are not limited to, cognitive and physical functioning ^26^, centenarian status ^25,27^, Down syndrome ^28^, HIV infection ^29^, obesity ^30^ and early menopause ^31^.

We demonstrated that the human pan-tissue clock can be directly applied to chimpanzee DNA methylation profiles ^20^, but its performance with profiles of other animals decline as a result of evolutionary genome sequence divergence. Recently, epigenetic clocks for mice were developed and used successfully to evaluate and confirm gold-standard longevity interventions such as calorie restriction and ablation of growth hormone receptor ^32–37^. These observations strongly suggest that age-related DNA methylation change is an evolutionarily conserved trait and as such, accurate age estimators as those developed for humans can be applied across species. Here we describe the development and performance of six different epigenetic clocks for rats. Two of these epigenetic clocks apply to both humans and rats. We used these six epigenetic clocks to evaluate the plasma fraction-based treatment in 4 rat tissues from 2-year-old rats. The corroborating results from these epigenetic clocks, together with physiological, histological and cognitive assessments demonstrate that administration of the plasma fraction reversed aging in rats.

## RESULTS

### DNA methylation data

All DNA methylation data were generated on a custom methylation array that applies to all mammals. We obtained in total, DNA methylation profiles of n=517 samples from 13 different tissues of rat (*Rattus norvegicus*) (**Supplementary Table 1, 2 and 3**), with ages that ranged from 0.0384 years (i.e. 2 weeks) to 2.3 years (i.e. 120 weeks). The rat tissue samples came from 3 different countries: (i) India (Nugenics Research in collaboration with NMIMS School of Pharmacy’s), (ii) the United States (H. Chen and L. Solberg Woods), and (iii) Argentina (R. Goya). Unsupervised hierarchical clustering shows that the methylation profiles clustered by tissue type, as would be expected (**Supplementary Figure 1**).

Our DNA methylation-based age estimators (epigenetic clocks) were developed (“trained” in the parlance of machine learning) using n=517 rat tissues. The two epigenetic clocks that apply to both species were developed by adding n=850 human tissue samples to the rat training set. Both rat and human tissues were profiled on the same methylation array platform (HorvathMammalMethylChip40) that focuses on 36,000 highly conserved CpGs (**Methods**).

### Epigenetic clocks

Our six different clocks for rats can be distinguished along several dimensions (tissue type, species, and measure of age). Some clocks apply to all tissues (pan-tissue clocks) while others are tailor-made for specific tissues/organs (brain, blood, liver). The rat pan-tissue clock was trained on all available tissues. The brain clock was trained using DNA samples extracted from whole brain, hippocampus, hypothalamus, neocortex, substantia nigra, cerebellum, and the pituitary gland. The liver and blood clock were trained using the liver and blood samples from the training set, respectively. While the four rat clocks (pan-tissue-, brain-, blood-, and liver clocks) apply only to rats, the human-rat clocks apply to both species. The two human-rat pan-tissue clocks are distinct, by way of measurement parameters. One estimates *absolute* age (in units of years), while the other estimates *relative* age, which is the ratio of chronological age to maximum lifespan; with values between 0 and 1. This ratio allows alignment and biologically meaningful comparison between species with very different lifespan (rat and human), which is not afforded by mere measurement of absolute age.

To arrive at unbiased estimates of the six epigenetic clocks, we used a) cross-validation of the training data and b) evaluation with an independent test data set. The cross-validation study reports unbiased estimates of the age correlation R (defined as Pearson correlation between the age estimate (DNAm age) and chronological age) as well as the median absolute error (**Figure 1**). The cross-validation estimates of the age correlations for all six clocks are higher than 0.9. The four rat clocks exhibited median absolute errors that range from 0.12 years (1.4 months) for the rat blood clock to 0.189 years (2.3 months) for the rat pan-tissue clock, **Figure 1A-D**). The human-rat clock for age generated an age correlation of R=0.99 when both species are analyzed together (**Figure 1E**) but is lower when the analysis is restricted to rat tissues alone (R=0.83, **Figure 1F**). In contrast, the human-rat clock for *relative age* exhibits high correlation regardless of whether the analysis is done with samples from both species (R=0.96, **Figure 1G**) or only with rat samples (R=0.92, **Figure 1H**). This demonstrates that relative age circumvents the skewing that is inherent when absolute age of species with very different lifespans are measured using a single formula. This is due in part to the unequal distribution of training data at the opposite ends of the age range.

**Figure 1:**
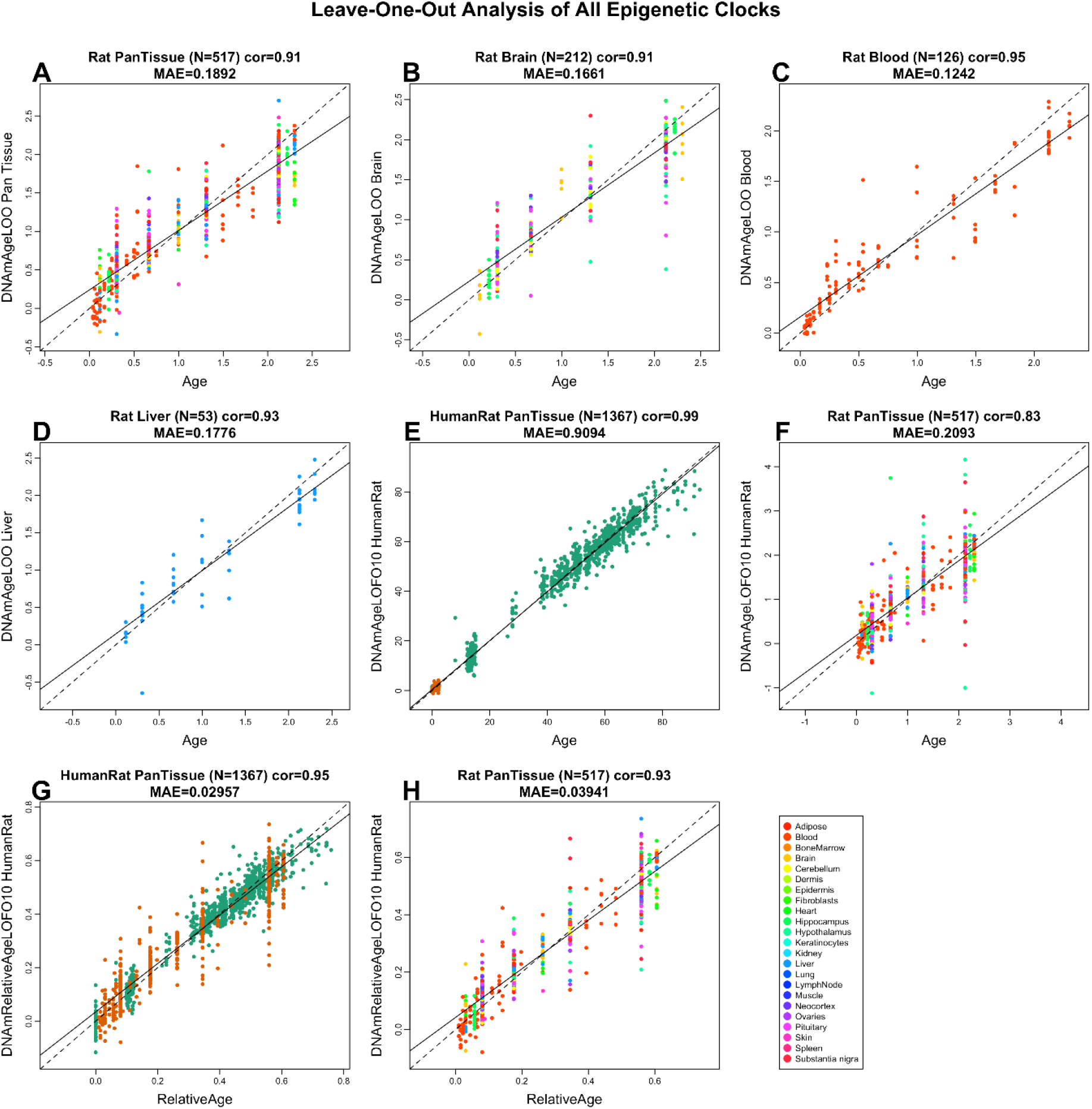
Cross-validation study of six epigenetic clocks for rat. A-D) Four epigenetic clocks that were trained on rat tissues only. E-H) Results for 2 clocks that were trained on both human and rat tissues. Leave-one-sample-out estimate of DNA methylation age (y-axis, in units of years) versus chronological age for A) Rat pan-tissue, B) Rat brain, C) Rat blood, and D) Rat liver clock. Dots are colored by A) tissue type or B) brain region. E and F) “Human-rat” clock estimate of absolute age. G, H) Human-rat clock estimate of relative age, which is the ratio of chronological age to the maximum lifespan of the respective species. Ten-fold cross-validation estimates of age (y-axis, in years) in E, G) Human (green) and rat (orange) samples and F, H) rat samples only (colored by tissue type). Each panel reports the sample size, correlation coefficient, median absolute error (MAE).

As indicated by its name, the rat pan-tissue clock is highly accurate in age estimation of all the different tissue samples tested and its performance with individual tissues can be more clearly seen in **Supplementary Figure 2**. We also evaluated the accuracy of the six epigenetic clocks in independent test data from the plasma fraction test study. In the untreated rat tissue samples, the epigenetic clocks exhibited high age correlations in all tissues (R>=0.95 in blood, liver, and the hypothalamus and R>=0.86 in Heart Tissue, **Supplementary Figure 3**).

### Rejuvenation effect of the plasma fraction treatment

The ability to generate epigenetic age clocks for specific rat organs can be readily appreciated from the perspective of their practical utility in aging research. However, the appreciation of a pan-tissue clock is much deeper as it extends into the conceptual aspect with regards to aging. The ability to estimate the age of different rat organs with a single clock; be it the rat pan-tissue clock or the human-rat pan-tissue clock, is a very strong indicator that epigenetic age is regulated across all tissues of an organism, and this regulation is mediated systemically. This in turn implies that it may be possible to centrally alter the rate of aging across different tissues of the body; a principle that underlies the plasma fraction treatment.

We applied the six clocks to an independent test data set (n=76) comprising four rat tissues (blood, liver, heart, hypothalamus). The main purpose of this is to test the hypothesis that the plasma fraction treatment reverses the epigenetic ages of 2-year-old rats.

Plasma fraction treatment was administered to rats following the experimental plan depicted in **Supplementary Figure 4.** Briefly, 18 Sprague Dawley rats were divided into three groups. A group of 6 young rats (30 weeks old), a second group of 6 old rats (109 weeks old) and a third group of 6 plasma fraction-treated old rats (also 109 weeks old). Plasma fraction treatment consists of two series of intravenous injections of plasma fraction. Rats were injected four times on alternate days for 8 days. A second identical series of injections were administered 95 days later. In its entirety, the experiment lasted 155 days. For the duration of the experiment, blood was drawn at regular intervals for haematological and biochemical analyses to monitor the impact of the treatment on blood, and solid vital organs. Cognitive functions of the rats were assessed four times during this period and at the end of the experiment, the animals were sacrificed, and DNA methylation profiles of several organs were generated. The DNA methylation profiles were evaluated using the above-mentioned six epigenetic clocks. The results derived from these profiles are plotted in **Figure 2,** which shows that the epigenetic ages of the old and young rats are readily distinguishable by all the six clocks. Crucially, plasma treatment of the old rats reduced the epigenetic ages of blood, liver and heart by a very large and significant margin, to levels that are comparable with the young rats. According to the six epigenetic clocks, the plasma fraction treatment rejuvenated liver by 73.4% (ranging from 63% to 81% depending on the clock, **Supplementary Table 8**), blood by 52% (ranging from 47 to 56%), heart by 52% (ranging from 40 to 74%), and hypothalamus by 11% (ranging from 1 to 20%). The rejuvenation effects are even more pronounced if we use the final versions of our epigenetic clocks: liver 75%, blood 66%, heart 57%, hypothalamus 19%. According to the final version of the epigenetic clocks, the average rejuvenation across four tissues was 54.2%. In other words, the treatment more than halved the epigenetic age (**Figure 2I-P**).

**Figure 2:**
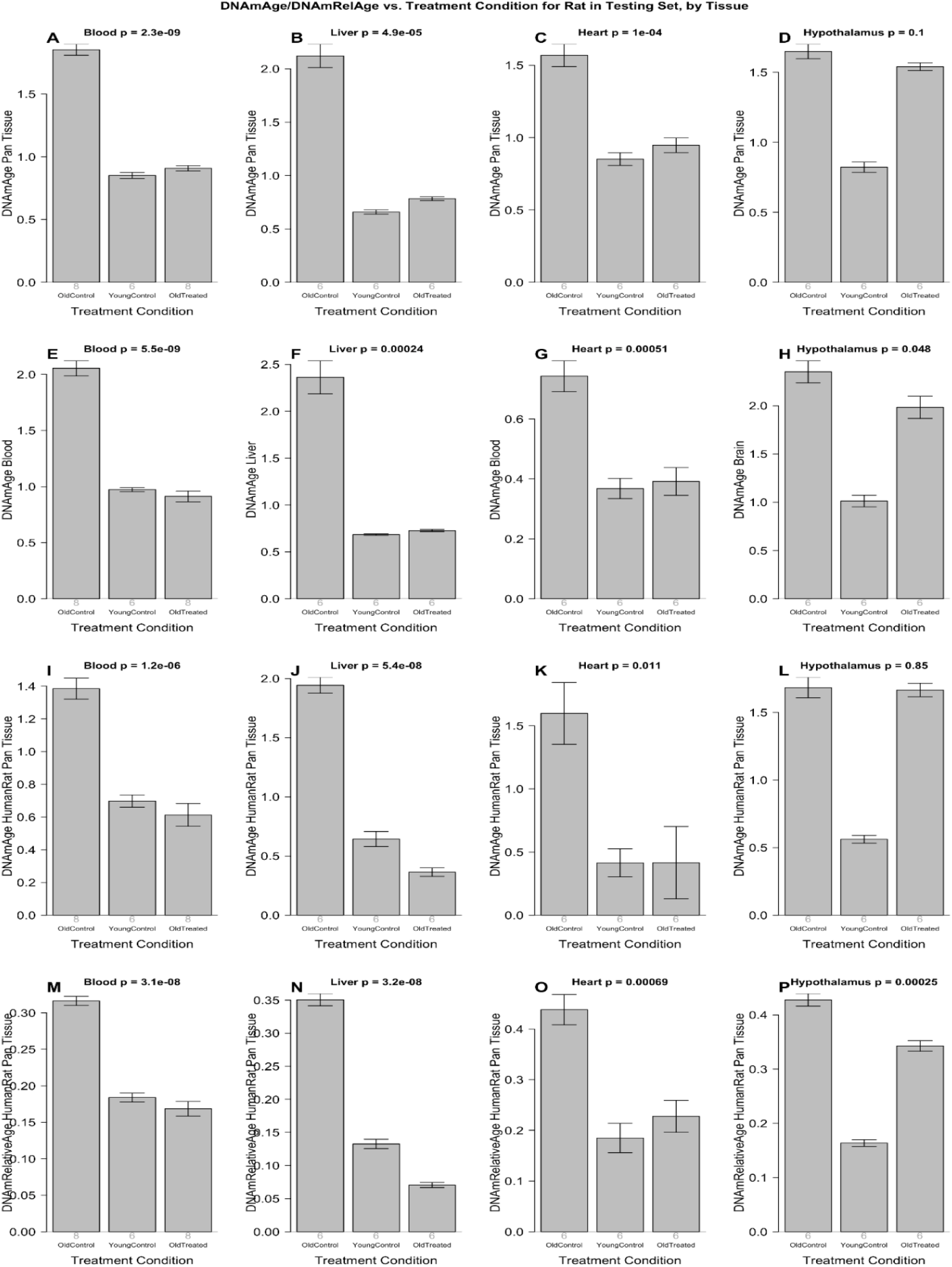
Epigenetic clock analysis of plasma fraction treatment. Six epigenetic clocks applied to independent test data from four rat tissue type (columns): blood, liver, heart, and hypothalamus. A-D) Rat pan-tissue clock. E) Rat blood clock applied to blood. F) Rat liver clock applied to liver. G) Rat blood clock applied to heart. H) Rat brain clock applied to hypothalamus. I-L) Human-rat clock measure of absolute age. M-P) Human-rat clock measure of relative age defined as age/maximum species lifespan. Each bar-plot reports the mean value and one standard error. P values results from analysis of variance. Student T-test p values result from a 2 group comparison of old controls (left bar) versus old treated samples (right bar), i.e. the young controls were omitted.

Epigenetic clocks are attractive aggregate biomarkers because they summarize the information of many CpGs into a single number (the age estimate). However, it can also be informative to look at individual CpGs in response to treatment. Meta-analysis of age-related methylation of individual CpGs across 13 untreated rat tissues, correlated negatively with that of 4 tissues from plasma fraction-treated animals (r=−0.62, **Supplementary Figure 5A**). This inverse correlation could also be observed when the tissues were individually analyzed instead. For blood, treatment-induced methylation changes correlated negatively, r=−0.64, with those that altered with age (**Supplementary Figure 5D**). For brain, liver and heart, the inverse correlations were r=−0.38 (**Supplementary Figure 5C**), r=−0.61 (**Supplementary Figure 5E**) and r=−0.47 (**Supplementary Figure 5F**) respectively. In summary, methylation at individual CpGs that normally alters with age was effectively reversed by plasma fraction treatment.

#### Physical and overt effects

The reduction of epigenetic age of plasma fraction-treated rats is particularly significant as it would appear to indicate that aging is a coordinated process as opposed to a stochastic one that occurs independently between the different organs. Before any further consideration of this notion, it is necessary to determine whether the reduction in epigenetic age is indeed biologically meaningful. In other words, is the rejuvenation of epigenetic age accompanied by changes in other well-characterized age-related endpoints. Equally important is the need to determine whether plasma fraction treatment generated any adverse side-effects.

The weight of rats for the duration of the experiment was monitored at regular intervals and the plots in **Supplementary Figure 6A** indicate that plasma fraction treatment did not affect food intake and appetite in any way, as the weight of the treated and untreated rats were similar, and there was no presentation of any overt signs of physical or behavioral abnormality. These features replicated those we observed in a mid-term (30-day) pilot experiment, where in addition, we also measured and found grip strength of old rats to be considerably improved by this treatment (**Supplementary Figure 6B**). At 15 days post-treatment, the strength of plasma fraction-treated old rats was indistinguishable from that of young ones. These and other encouraging results from the mid-term pilot studies, prompted a longer-term (155-day) investigation, with a new preparation of plasma fraction. The results from this study forms the main corpus of this report. These two independent investigations produced similar results that varied only in terms of magnitude, as is consistent with their different durations. Histological examinations of the various organs did not indicate any obvious abnormalities after 155 days of treatment (**Supplementary Figure 7** and **Supplementary Table 4**). Instead, oil red O staining showed that accumulation of fat in old tissues was greatly reduced in plasma fraction-treated rats (**Supplementary Figure 8**).

#### Blood cell indices and haematology

To monitor potential effects of plasma fraction on the blood of rats, we measured haemoglobin levels, mean corpuscular volume (MCV), mean corpuscular haemoglobin (MCH), mean corpuscular haemoglobin concentration (MCHC), hematocrit (HCT) levels and obtained counts of red blood cells, platelets, white blood cells and lymphocytes at 0, 60 and 155 days from the start of treatment. These blood indices are informative indicators of malfunction of bone marrow and vital organs and importantly, they vary with the age of the animal. In **Supplementary Figure 9,** it can be seen that all these blood indices were very different between young and old rats at the start of the experiment, and in time, plasma fraction treatment nudged all these parameters, with the exception of platelets, away from the values exhibited by old untreated rats towards those of the young ones. This is easily visualized by the movement of the orange dots, representing treated old rats in each graph towards the blue dots, representing young rats. Plasma fraction treatment has not caused any changes to the blood indices that would indicate any organ dysfunction. Instead it rejuvenated the blood of the rats, which is consistent with the significant reduction of epigenetic age of their blood as measured by both the rat multi-tissue and blood clocks.

#### Biomarkers for vital organs

To ascertain the impact of plasma fraction treatment on vital organs, we measured the levels of the following biomarkers on 30, 60, 90, 120 and 155 days from the start of the experiment: bilirubin, serum glutamic-pyruvic transaminase (SGPT) and serum glutamic-oxaloacetic transaminase (SGOT) to monitor liver function; triglycerides (TG), HLD and cholesterol to monitor risk of atherosclerosis and heart disease, and liver function as well; glucose to monitor the pancreas and diabetes; and creatinine and blood urea nitrogen for kidney function. The levels of all these biomarkers in the treated old rats were altered towards the values of young rats, without exception **(Figure 3)**. This is easily visualized by the movement of the orange dots, representing treated old rats in each graph towards the blue dots, which represent young rats. Collectively, these results show that the function of all the vital organs tested through their respective biomarkers, were rejuvenated by plasma fraction treatment. This is entirely consistent with reversal of the epigenetic ages of their hearts and livers.

**Figure 3:**
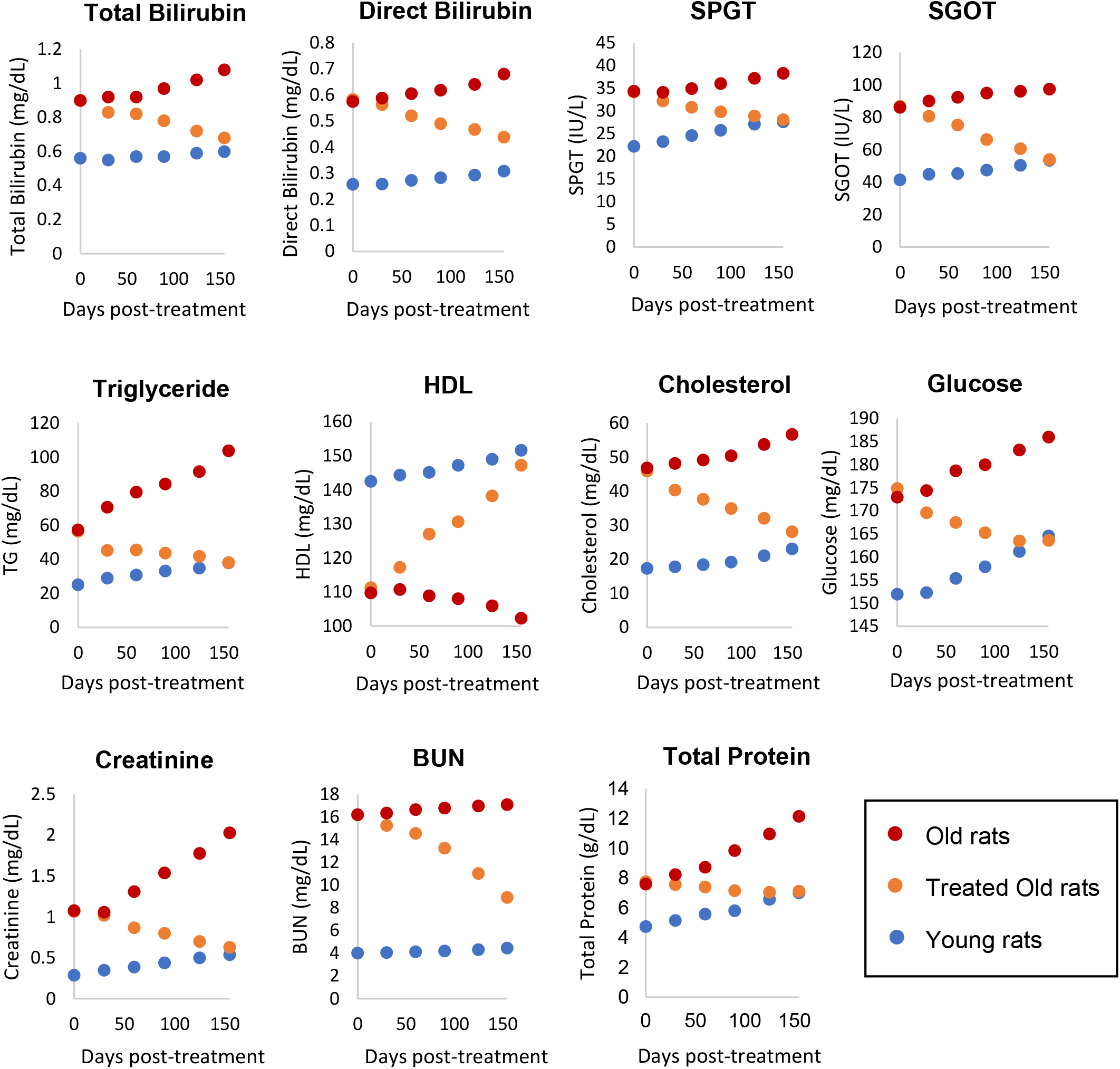
Assessment of plasma fraction treatment on vital organ functions. At 0, 30, 60, 90, 120 and 155 days from commencement of the experiment, the health and function of vital organs (liver, heart, kidney and pancreas) of 6 rats per group were monitored through measurements of appropriate biomarkers. SPGT = serum glutamic-pyruvic transaminase, SGOT = serum glutamic-oxaloacetic transaminase, HDL = high-density lipoprotein and BUN = blood urea nitrogen. Red dots represent data points of old rats, orange dots represent treated old rats and blue represent young rats. For clarity, the plotted data points represent average values from 6 rats each. Detailed measurements of each parameter with standard deviations are provided in Supplementary table 7.

#### Cognitive function

The decline of cognitive function is a well-characterized feature of increasing age. Learning and memory, which are constituent characteristics of cognitive functions, decline not only in human but also in rats, starting from 12 months of age 38. Barnes maze was used to measure the latency period required by the rats to escape through the right hole into an escape box. Videos illustrating the latency pattern of three rats (young, old treated, old untreated) can be found in the Supplement. Within a month of plasma fraction treatment, the rats exhibited significantly reduced latency to escape (**Figure 4**), i.e., they learned and remembered better. After the second month, the treated rats began with a slightly reduced latency period compared to the untreated old rats, and once again, they learned much faster than the latter. By the third month, it was clear that treated rats remembered the maze much better than the untreated ones even from the first day of test as their latency period was significantly reduced and by the end of the test period their latency was similar to that of the young rats. This feature was sustained and repeated in the fourth month. Collectively, these results show that plasma fraction improved the learning and memory of the rats. Interestingly, the epigenetic age of treated rat brain samples (hypothalamus) was lower than the untreated old ones, but less markedly than the magnitude of decrease of epigenetic age of the blood, heart and liver. This introduces numerous possible insights into the relationship between cognitive function, biological age and physical health which will be further elaborated in the discussion.

**Figure 4:**
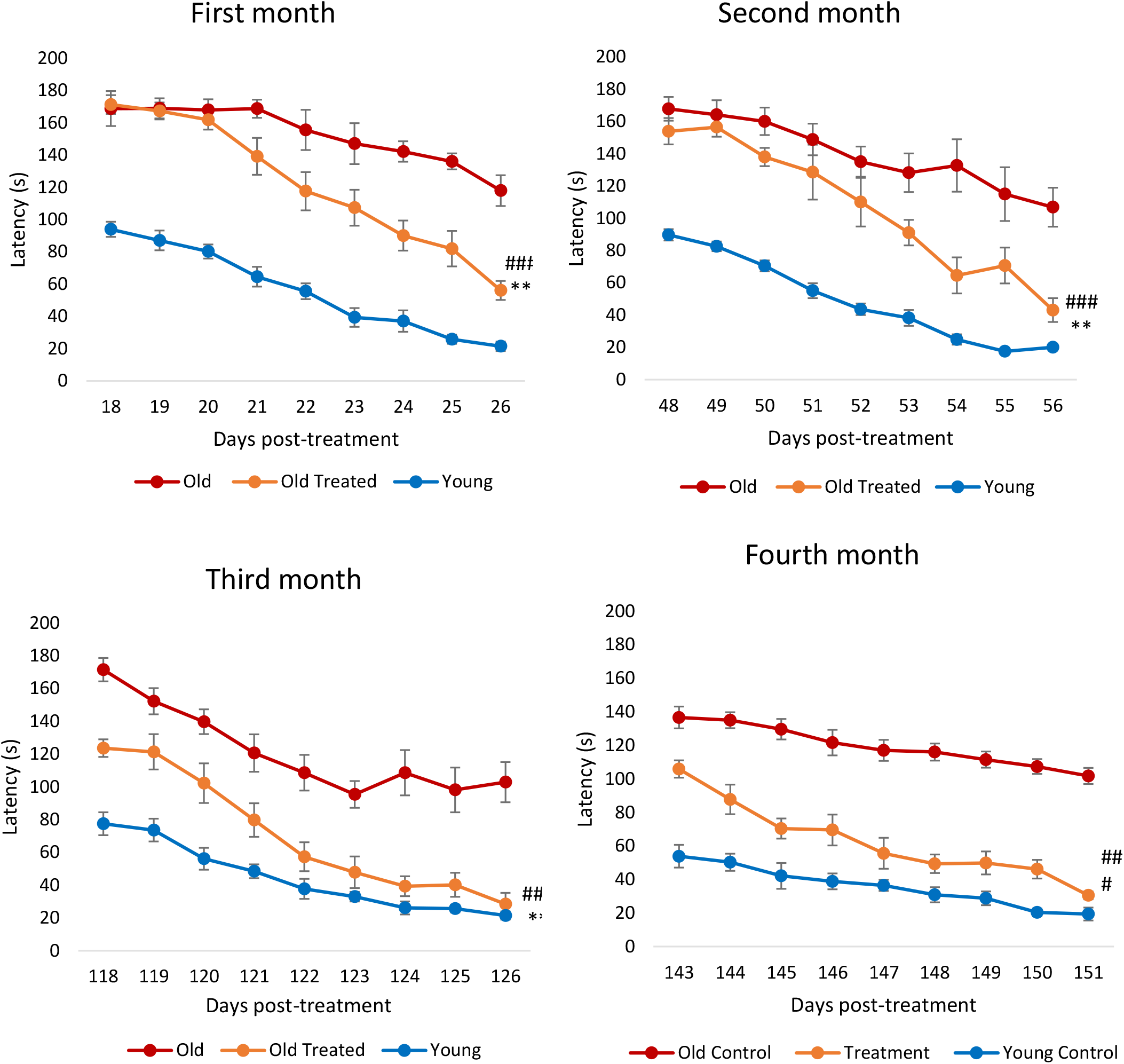
Assessment of plasma fraction treatment on cognitive function (learning and memory). Rats were subjected to Barnes maze test in the first to fourth month from commencement of the experiment. Each assessment consists of nine consecutive days of test where the time (in seconds) required by the rats to find the escape hole (latency) was recorded and plotted. The error bars depict 2 standard errors.

#### Cellular stress

There is no doubt that the reduction in epigenetic age of liver, heart, brain and blood by plasma fraction was accompanied by startling improvement in the function of these organs. In addition to the decline of organ function with age, is the rise of two cell stress features which are related, namely oxidative stress and chronic inflammation; the excess of which have been linked to multiple pathologies. We carried out a panel of biomarker tests for these two features in the test rats.

#### Oxidative stress

Oxidative stress results from the excess of reactive oxygen species (ROS) in cells. This situation can arise due to the over-production of ROS or the decline in the ability to remove or neutralize ROS. While ROS at low levels are not harmful and are even necessary, at higher levels, they interact with biomolecules such as lipids and compromise their function. Measuring the levels of malondialdehyde (MDA), which is the end-product of poly-unsaturated fatty acid peroxidation, reveals the levels of cellular ROS. The amounts of MDA were clearly higher in the brain, heart, lung and liver of older rats (**Figure 5**), and plasma fraction treatment reduced this level to that of young rats. Hence, regardless of the source of augmented ROS in older rats, plasma fraction appears to be able to reduce it effectively.

**Figure 5:**
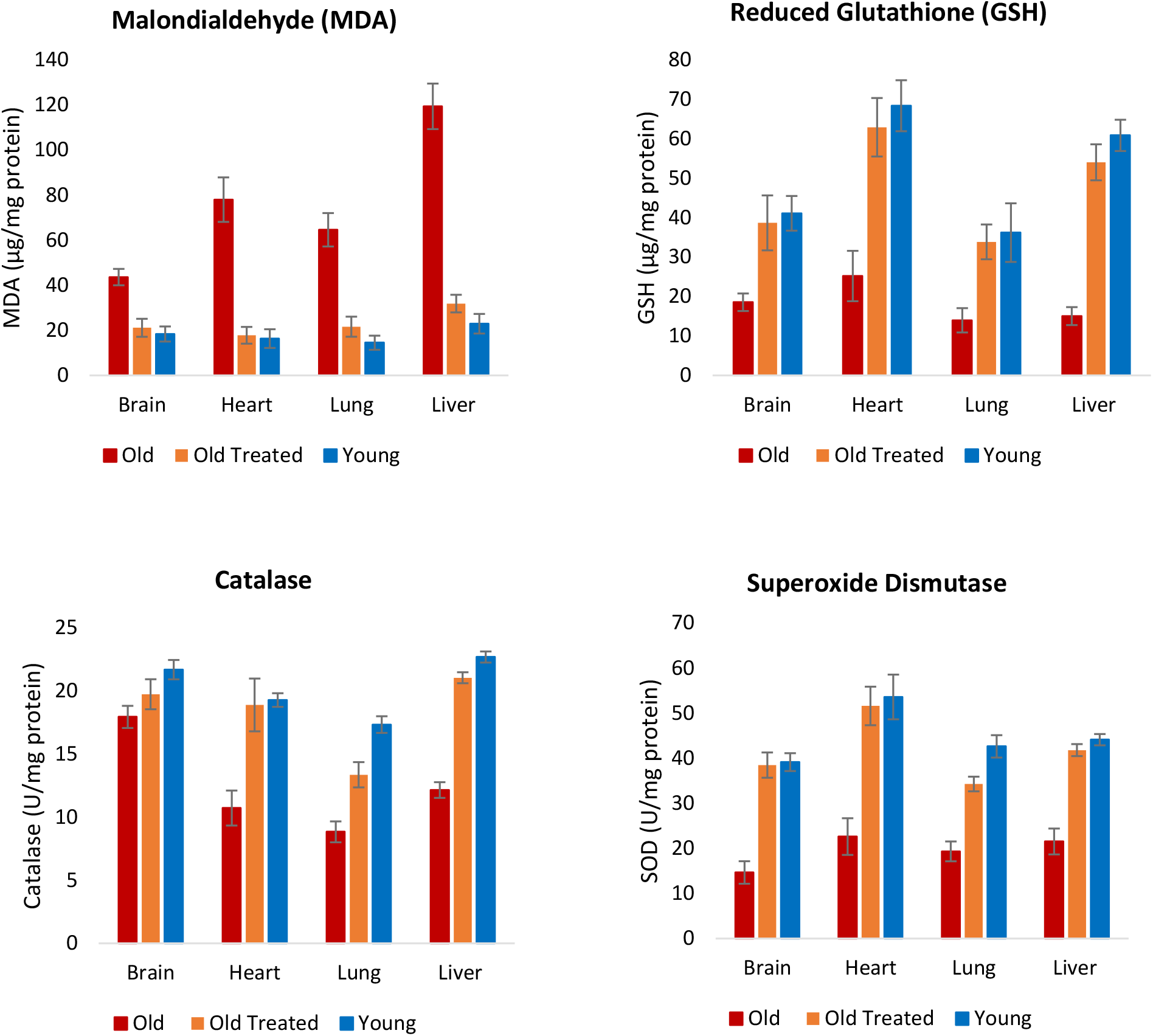
Assessment of plasma fraction treatment on oxidative stress. At the end of the experimental period of 155 days, lipid peroxidation level, which is an indicator of intracellular level of reactive oxygen species (ROS) was determined by measuring the amount of Malondialdehyde (MDA), which is the end product of polyunsaturated fatty acid peroxidation. The levels of three anti-oxidants; reduced glutathione, catalase and superoxide dismutase levels were also measured to ascertain the impact of plasma fraction treatment on oxidative stress. The error bars depict 2 standard errors.

Apart from increased production of ROS, decreased efficiency in eliminating ROS also contributes to the age-associated rise in its level. ROS are neutralized by cellular anti-oxidants including but not limited to reduced glutathione, catalase and superoxide dismutase, which all work in very different ways. The levels of these three anti-oxidants were reduced in tissues of old untreated rats but the plasma fraction treatment augmented them to levels that are comparable to young ones (**Figure 5**).

It remains to be ascertained if the reduction of ROS levels, as measured by MDA, was due entirely to the increase in the amounts of anti-oxidants, or whether plasma fraction treatment also induced a concomitant reduction in the production of ROS from multiple and various intracellular sources. Nevertheless, the end-point of ROS levels in old rats being diminished to the level of young ones is yet another indication of the rejuvenating effect of the treatment.

#### Chronic inflammation

Inflammation is an important response that helps protect the body, but excess inflammation especially in terms of duration of this response can have very detrimental effects instead. This occurs when inflammation fails to subside and persists indefinitely; a condition referred to as chronic inflammation, which for reasons not well-understood, increases with age and is associated with a multitude of conditions and pathologies. The levels of two of the most reliable and common biomarkers of chronic inflammation; interleukin 6 (IL-6) and tumor necrosis factor α (TNF-α) are found to be considerably higher in old rats (**Figure 6**), and these were very rapidly diminished, within days by plasma fraction treatment, to comparable levels with those of young rats. This was especially stark with IL-6. In time, the levels of these inflammatory factors began to rise gradually, but they were once again very effectively reduced following the second administration of the plasma fraction on the 95^th^ day. The reduction of these inflammation markers is consistent with the profile of the nuclear factor erythroid 2-like 2 protein (Nrf2), which plays a major role in resolving inflammation, in part by inhibiting the expression of IL-6 and TNF-α. Nrf2 also induces the expression of antioxidants that neutralizes ROS, which is also a significant feature in inflammation ^39^. In summary, plasma fraction reduces oxidative stress and chronic inflammation, which are age-associated pan-tissue stresses, to the levels found in young rats.

**Figure 6:**
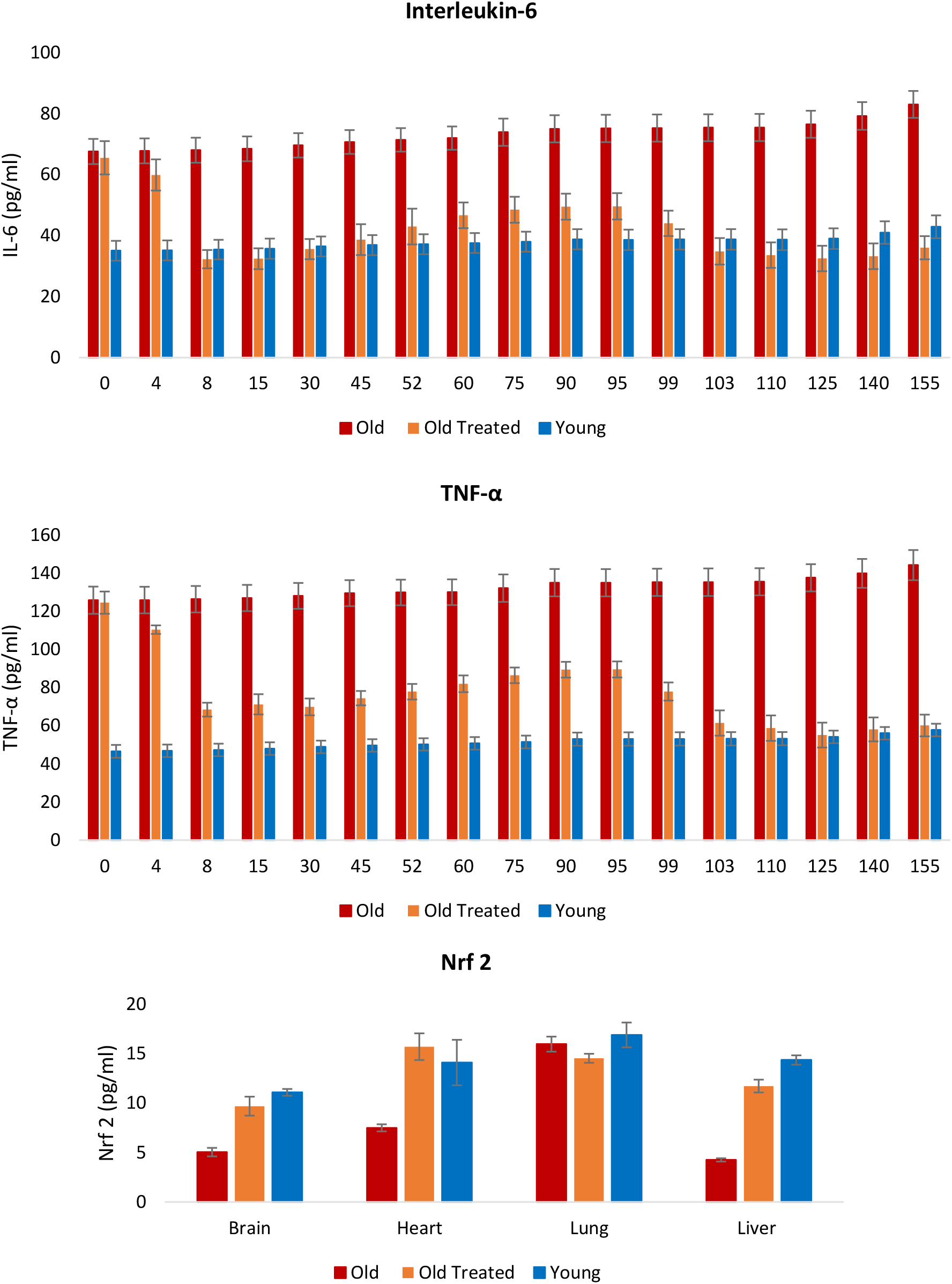
Assessment of the effect of plasma fraction treatment on chronic and systemic inflammation. Blood levels of interleukin-6 (IL-6) and tumor necrosis factor α (TNF-α) were measured at regular intervals throughout the 155-day period of the experiment. At the end of the experiment the levels of Nrf2, a pivotal modulator of inflammation and oxidative stress were measured in brain, heart, lung and liver of the rats. The error bars depict 2 standard errors.

#### Cellular senescence

One of the best characterized contributors to aging is the senescent cell. Cells become senescent due to numerous possible causes including exhaustive replication (replicative senescence), over-expression of oncogene or chronic DNA damage signaling owed to un-repaired DNA. Many senescent cells switch on the expression of acidic beta-galactosidase, which is known as senescence-associated beta-galactosidase (SA-β-galactosidase). As the presence of this enzyme activity signals the senescent state of cells, SA-β-galactosidase is used as a biomarker of senescent cells. Senescent cells are stained blue when provided with SA-β-galactosidase substrate in acidic pH, as seen in high levels in the brains and livers of old rats (**Figure 7**). Plasma fraction treatment reduced the level of senescent cells by a very considerable degree. The heart and lung of old rats possessed very few senescent cells, and this did not change after treatment.

**Figure 7:**
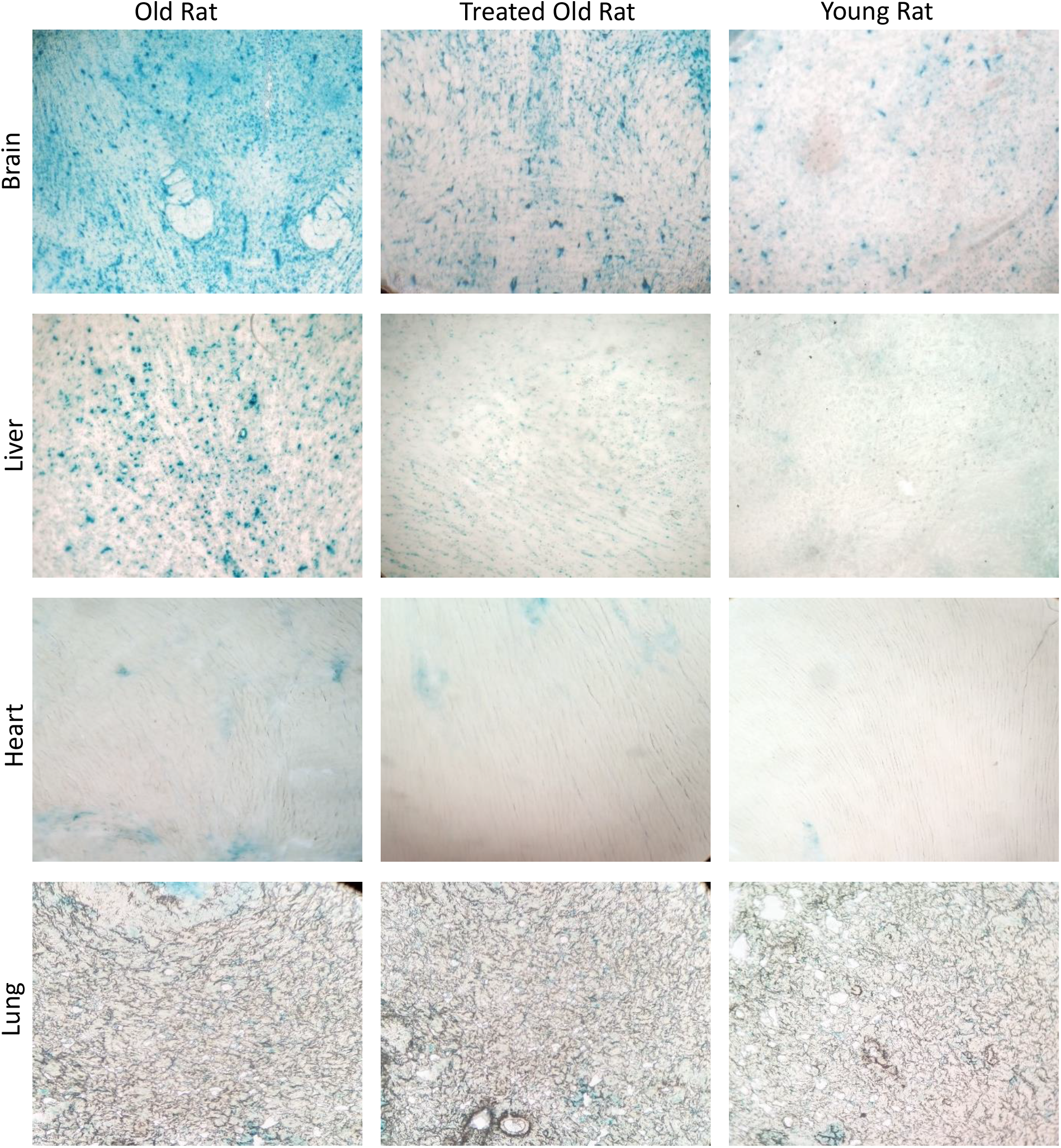
Assessment of the effect of plasma fraction treatment on cellular senescence. At the end of the experimental period of 155 days, senescent cells in brain, heart, lung and liver of old rats, plasma fraction-treated old rats and young rats were stained for senescence-associated beta-galactosidase, whose activity on the substrate provided turns senescent cells blue in color.

## Discussion

This study consists of two connected parts; the first of which concerns the development of six rat epigenetic clocks, which were subsequently used in the second part to interrogate the effects of plasma fraction treatment on age.

### Development of rat epigenetic clocks

Epigenetic clocks for humans have found many biomedical applications including the measure of age in human clinical trials ^22,40^. These clocks provide a standard measure of DNA methylation state in function of chronological age. As impressive as its accuracy is, it is the divergence from this standard that was particularly important because it uncovered the association between accelerated epigenetic age and the associated increased risk of a host of conditions and pathologies, indicating that epigenetic clocks are associated with biological age. This instigated development of similar clocks for animals, of which the ones for mice were particularly attractive as they allow for epigenetic age to be modeled in a mouse system, and at the same time allows existing mouse models of aging to be interrogated with regards to epigenetic aging. Indeed, numerous mouse epigenetic clocks have since been developed and successfully validated against factors, such as rapamycin, caloric restriction and growth factor ablation, which are all well-characterized in their effects on aging of mice ^32–37^. While the advantages of mouse as a biological model lies in no small part to their size, this also poses a limitation in studies that require regular interval collection of sufficient amounts of blood for analyses, as was the case in the second part of this study. The development of six rat epigenetic clocks described here was based on novel DNA methylation data that were derived from thirteen rat tissue types. The two human-rat clocks demonstrate the feasibility of building epigenetic clocks for two species based on a single mathematical formula. A critical step toward crossing the species barrier was the use of a mammalian DNA methylation array that profiled 36 thousand probes that were highly conserved across numerous mammalian species. The rat DNA methylation profiles represent the most comprehensive dataset thus far of matched single base resolution methylomes in rats across multiple tissues and ages. We expect that the availability of these clocks and their impressive performance in the second part of this study will provide a significant boost to the attractiveness of the rat as biological model in aging research.

Beyond their utility, these clocks reveal several salient features with regards to the biology of aging. First, the rat pan-tissue clock re-affirms the implication of the human pan-tissue clock, which is that aging might be a coordinated biological process that is harmonized throughout the body. Given that the circulatory system irrigates and connects all the organs, it is more likely than not, that the regulation and harmonization of age are mediated systemically. Second, the ability to combine these two pan-tissue clocks into a single human-rat pan-tissue clock attests to the high conservation of the aging process across two evolutionary distant species. This implies, albeit does not guarantee, that treatments that alter the epigenetic age of rats, as measured using the human-rat clock is likely to exert similar effects in humans. If validated, this would be a step change in aging research. Although conservation of aging mechanism could be equally *deduced* from the existence of multiple individual clocks for other mammals (mouse, dog), the single formula of the human-rat clock that is equally applicable to both species effectively *demonstrates* this fact. It is evident that the mechanism underpinning aging is a very primitive and important biological process, which ensured its conservation across the mammalian kingdom through time.

The incorporation of two species with very different lifespans such as rat and human, raises the inevitable challenge of unequal distribution of data points along the age range. The clustering of the shorter and longer lifespan species at the lower and higher age range respectively can skew the accuracy of the clock when it is applied individually to either of the species. This effect is mitigated by the generation of the human-rat pan-tissue *relative age* clock which embeds the estimated age in context of the maximal lifespan recorded for the of the relevant species. In addition to minimizing the skew, this mathematical operation also generates a much more biologically meaningful value because it indicates the relative biological age and fitness of the organism in relation to its own species. This principle will be an important feature to incorporate in future composite epigenetic clocks of different species.

### Plasma fraction treatment

Although parabiosis experiments showing organs of old mice to benefit biologically from sharing circulation with young ones were carried out decades ago, little follow-up work was done to capitalize on this remarkable observation until recently, when these experiments were successfully repeated and greatly expanded ^1–8,10^. Young plasma transfer was shown to also work ^8^. This was important for two reasons, the first being that it confirmed the notion that it is indeed blood factors that were improving the old animals and not to them benefiting from the healthier and better functioning vital organs of the young animals. Second, it provided a far less demanding and more humane way to investigate this phenomenon.

The improvement of organ function of old animals was rapidly and intuitively assumed to indicate rejuvenation. This, however, is not the only valid interpretation, as it is equally possible that the organs of the old animals were improved in function, but with no impact on their epigenetic state or epigenetic age. It raised the perennial question of what is aging. If aging is defined purely in functional terms as decline in organ and tissue function, then improvement of these functions could logically be interpreted as rejuvenation. It is evident that this perspective would encounter numerous conceptual obstacles, in real-life and theoretical scenarios, which will not be elaborated here. This problem would have remained intractable if age could only be measured by time and not by a molecularly-based method.

The improvements in the function of all the organs tested correlated with a substantial reduction in their epigenetic ages, with the exception of the brain, where the reduction is significant for only some of our rat clocks (e.g. the brain clock and the human-rat clock of relative age). The fact that the rats perform better in the maze test has two non-exclusive explanations: it may be a consequence of a molecular rejuvenation of the brain or that it reflects healthier brain function due to overall physical improvements.

This, of course, raises a related and equally interesting question, which is why does plasma fraction treatment not reduce brain epigenetic age by the same magnitude as it does the other organs? We can only begin to address this question after having first understood what epigenetic aging entails. As it stands, our knowledge in this area remains limited, but it is nevertheless clear that: (a) epigenetic aging is distinct from the process of cellular senescence and telomere attrition ^41^, (b) several types of tissue stem cells are epigenetically younger than non-stem cells of the same tissue ^42,43^, (c) a considerable number of age-related methylation sites, including some clock CpGs, are proximal to genes whose proteins are involved in the process of development ^44^, (d) epigenetic clocks are associated with developmental timing ^22,45^, and (e) relate to an epigenomic maintenance system ^20,46^. Collectively, these features indicate that epigenetic aging is intimately associated with the process of development and homeostatic maintenance of the body post-maturity. While most organs of the body turn-over during the lifetime of the host, albeit at different rates, the brain appears at best to do this at a very much slower rate ^27,47^. While most tissues harbor stem cells that are necessary for replenishment and turnover, stem cells in adult brain have only been detected in a defined and very limited area of the subventricular zone, olfactory bulb (in rats), hippocampus and hypothalamic proliferative region ^48^. As such, if plasma fraction treatment’s rejuvenating effect is mediated through the process of development and involves tissue stem cells, then its effect on the epigenetic age of the brain would appear to be modest, which indeed it does. It is to be noted however, that improving brain function does not depend on neurogenesis as much as it does on synapse formation and factors such as NMDA receptors which decline in density with age ^49,50^.

Apart from rejuvenating the vital organs of the treated rats, plasma fraction also impacted two fundamental physiological processes that underlie a great number of pathologies, namely oxidative stress and inflammation. Within a week of treatment, the markers of chronic inflammation (IL-6 and TNF-α) were significantly reduced and remained low throughout the entire experiment. Likewise, markers of oxidative stress in brain, heart, lung and liver, which were very much higher in control old rats, were at the end of the experimental period, indistinguishable between plasma fraction-treated old rats and young ones. Concomitant with this drastic reduction in oxidative stress was the augmented levels of anti-oxidants (GSH, Catalase and SOD) in these tissues, indicating that modulating the levels of ROS to that of youthful rats is at least one way by which plasma fraction suppresses oxidative stress. It remains to be ascertained whether the rate of ROS generation is also reduced. The levels of Nrf2, a transcription factor that impacts on oxidative stress, as well as inflammation, were raised by plasma fraction treatment of old rats to those of the young ones, indicating yet another level by which this treatment modulates these two critical processes. Collectively, these results show that plasma fraction treatment impacts not only the overt performances of organs, but also the underlying physiological processes that are pivotal for optimal organ function and health.

The plasma fraction treatment system employed here has been tested several times with reproducible success. In addition to improvements in clinical markers of aging, our work shows, for the first time, that plasma fraction treatment also reduces considerably, the epigenetic age of multiple tissues. It is evidently very effective in rejuvenating several tissues in old rats; requiring only two series of injections. The rejuvenation of non-blood organs from this intravenous treatment, coupled with reversal of the epigenetic clock within these organs supports the notion that aging can be systemically controlled, at least in part through the circulatory system with plasma as the medium. It is important to appreciate the potential step change in health care that plasma fraction treatment and epigenetic clocks can introduce. Instead of treatment of one disease at a time and eventually co-morbidities, plasma fraction treatment, by rejuvenation of the body, may be able to systemically reduce the risk of the onset of several diseases in the first place.

## Materials and Methods

### Materials

In total, we analyzed n=593 rat tissue samples from 13 different sources of DNA (**Supplementary Table 1**). Ages ranged from 0.0384 years (i.e. 2 weeks) to 2.3 years (i.e. 120 weeks). The rat data were comprised of a training set (n=517) and a test set (n=76). To build human-rat clocks, we added n=850 human tissue samples to the training data. We first trained/developed epigenetic clocks using the training data (n=517 tissues). Next, we evaluated the data in independent test data (n=76 for evaluating the effect of plasma fraction treatment (**Supplementary Table 3**). We used n=517 tissue to train 4 clocks: a pan-tissue clock based on all available tissues, a brain clock based on regions of the whole brain - hippocampus, hypothalamus, neocortex, substantia nigra, cerebellum, and the pituitary gland, a liver clock based on all liver samples, and a blood clock.

### Methods

The rat tissues came from 4 different labs across three countries:(i) India: Nugenics Research in collaboration with School of Pharmacy SVKM’s NMIMS University (K. Singh), (ii) United States: University of Tennessee Health Science Center (H. Chen) and Medical College of Wisconsin (L.C. Solberg Woods), and (iii) Argentina: University of La Plata (R. Goya).

### Rats from Tennessee and Wisconsin

Blood samples (n=48): Male and female heterogeneous stock rats were bred at the Medical College of Wisconsin (Solberg Woods Lab) or University of Tennessee Health Science Center (Hao Chen Lab). Heterogeneous Stock (HS) populations were originally developed by breeding together eight inbred strains, followed by maintaining the colony in a manner that minimizes inbreeding, allowing fine-resolution genetic mapping of a variety of complex traits ^51^. Rats were euthanized at different ages by an overdose of isoflurane (> 5%). Trunk blood was collected immediately and stored at −80 °C until processing. Blood samples were treated with streptokinase (60-80 IU/200 μl blood, overnight incubation at 37 °C) and DNA was extracted using the QiaAmp Blood Mini Kit (Qiagen Cat No./ID: 51304) following manufacturer’s instructions. All procedures were approved by the Institutional Animal Care and Use Committee of the University of Tennessee Health Science Center or the Medical College of Wisconsin and followed the NIH Guide for the Care and Use of Laboratory Animals. Genomic DNA was isolated from tissue samples mostly using Puregene chemistry (Qiagen). DNA from liver was extracted manually and from blood using an automated Autopure LS system (Qiagen). From tissues and clotted blood samples DNA was extracted manually using QiaAmp DNA Blood Midi Kit and the DNeasy Tissue Kit according to manufacturer’s protocol (Qiagen, Valencia, CA). DNA from BA10 was extracted on an automated nucleic acid extraction platform Anaprep (Biochain) using a magnetic bead-based extraction method and Tissue DNA Extraction Kit (AnaPrep).

### Rats from the University of La Plata (R. Goya lab)

Multiple tissues/cell types (adipose, blood, cerebellum, hippocampus, hypothalamus, liver, neocortex, ovaries, pituitary, skin, substantia nigra): Young (3.7 mo., n=11), Late Adults (LA, 8.0 mo., n=9), Middle-Aged (M-A, 15.7 mo., n=6) and Old (25.5 mo., n=14) female Sprague-Dawley (SD) rats, raised in our Institute, were used. Animals were housed in a temperature-controlled room (22 ± 2ºC) on a 12:12 h light/dark cycle. Food and water were available *ad libitum*. All experiments with animals were performed in accordance to the Animal Welfare Guidelines of NIH (INIBIOLP’s Animal Welfare Assurance No A5647-01) and approved by our Institutional IACUC (Protocol # P05-02-2017).

#### Tissue sample collection

Before sacrifice by decapitation, rats were weighed, blood was withdrawn from the tail veins with the animals under isoflurane anesthesia and collected in tubes containing 10μl EDTA 0.342 mol/l for 500μl blood. The brain was removed carefully severing the optic and trigeminal nerves and the pituitary stalk (not to tear the pituitary gland), weighed and placed on a cold plate. All brain regions were dissected by a single experimenter (see below). The skull was handed over to a second experimenter in charge of dissecting and weighing the adenohypophysis. The rest of the body was handed to other 2 or 3 experimenters who dissected and collected whole ovaries, a sample of liver tissue, adipose tissue and skin tissue from the distal portion of tails.

#### Brain region dissection

Prefrontal cortex, hippocampus, hypothalamus, substantia nigra and cerebellum were rapidly dissected on a cold platform to avoid tissue degradation. After dissection, each tissue sample was immediately placed in a 1.5ml tube and momentarily immersed in liquid nitrogen. The brain dissection protocol was as follows. First a frontal coronal cut was made to discard the olfactory bulb, then the cerebellum was detached from the brain and from the medulla oblongata using forceps. To isolate the medial basal hypothalamus (MBH), brains were placed ventral side up and a second coronal cut was made at the center of the median eminence (−3,6 mm referred to bregma). Part of the MBH was taken from the anterior block of the brain and the other part from the posterior block in both cases employing forceps. The hippocampus was dissected from cortex in both hemispheres using forceps. This procedure was also performed on the anterior and posterior blocks, alternatively placing the brain caudal side up and rostral side up. To dissect the substantia nigra, in each hemisphere a 1-mm thick section of tissue was removed from the posterior part of the brain (−4,6 mm referred to bregma) using forceps. Finally, the anterior block was placed dorsal side up, to separate prefrontal cortex. With a sharp scalpel, a cut was made 2 mm from the longitudinal fissure, and another cut was made 5 mm from it. Additionally, two perpendicular cuts were made, 3 mm and 6 mm from the most rostral point, obtaining a 9 mm^2^ block of prefrontal cortex. This procedure was performed in both hemispheres and the two prefrontal regions collected in a code-labeled tube.

#### Anterior pituitary

Using forceps the dura matter that covers gland was removed leaving the organ free on the sella turcica. The neural lobe was carefully separated from the anterior pituitary (AP) which was then carefully lifted with fine curved tip forceps pointing upwards. It was rapidly weighed, then put in a tube and placed momentarily in liquid nitrogen.

#### Ovaries

The genital apparatus was dissected by cutting the mesentery to isolate the uterine horns, the tubular oviduct, the ovaries and the junction between the anus/rectum and the vulva/vagina, leaving the unit of the sexual organs and the urinary bladder isolated. The ovaries were carefully separated from the oviducts; the fat around the ovaries was also removed. Both gonads were placed in a single eppendorf tube and momentarily placed in liquid nitrogen.

#### Liver

Liver tissue extraction was made by cutting a piece of the median lobe (0.5 cm x 0.5 cm). Tissue was placed in a tube and momentarily stored immersed in liquid nitrogen.

#### Adipose tissue

Adipose tissue samples were obtained from the fatty tissue of the small intestine.

#### Tail skin

For skin tissue, 5 cm of a distal tail portion were cut with scissors. Skin was separated and hair removed using scalpel. Tissue was placed in a tube and stored as described for other tissues.

DNA was extracted from blood on an automated nucleic acid extraction platform called QiaSymphony (Qiagen) with a magnetic bead-based extraction kit, QIAsymphony DNA Midi Kit (Qiagen). DNA was extracted from tissue on an automated nucleic acid extraction platform called Anaprep (Biochain) with a magnetic bead-based extraction kit, Tissue DNA Extraction Kit (Biochain). DNA from brain regions was extracted using an automated nucleic acid extraction platform called QIAcube HT (Qiagen) with a column-based extraction kit, QIAamp 96 DNA QIAcube HT Kit (Qiagen).

### Rats from Nugenics Research Group

Sprague Dawley rats of both sexes were used, from which blood, whole brain, heart and liver were harvested. Two batches or samples were prepared: the first batch was intended for training the epigenetic clock: n=42 blood samples, n=18 whole brain, n=18 heart, n=18 liver samples. The second batch of test set involved n=76 tissue samples (n=22 blood, n=18 liver, n=18 heart, n=18 hypothalamus). The test data were used to evaluate the effect of the treatment in 3 conditions: young (30-week old) and treated old samples (109 weeks old), and untreated old samples (again 109 weeks old). We evaluated 4 sources of DNA: blood (n=18), liver (n=18), heart (n=18) and hypothalamus (n=18). Ethics committee approval number - CPCSEA/IAEC/P-6/2018.

Male Sprague Dawley rats of 8 weeks (200–250 g) and 20 months (400-450g) were procured from the National Institute of Bioscience, Pune, India. Animals were housed in the animal house facility of School of Pharmacy, SVKM’s NMIMS University, Mumbai during the study under standard conditions (12:12 h light: dark cycles, 55-70% of relative humidity) at 22±2°C temperature with free access to water and standard pellet feed (Nutrimix Std-1020, Nutrivet Life Sciences, India). The animals were acclimatized to laboratory environment for seven days before initiation of the study. The experimental protocol was approved by the Institutional Animal Ethics Committee. The approval number is CPCSEA/IAEC/P-75/2018. Rats were euthanized at different ages by an overdose of isoflurane (> 5%). Trunk blood was collected immediately and stored at −80 °C until processing. 100 μl blood sample was treated with 20 μl Proteinase K and then the volume was adjusted to 220 μl with Phosphate Buffer Saline (PBS) in 1.5 ml or 2 ml microcentrifuge tube. 200 μl buffer AL was mixed thoroughly to this mixture by vortexing, and incubated at 56°C for 10 min. Then 200 μl ethanol (96–100%) was added to the sample and mix thoroughly by vortexing and DNA were extracted using the Qiagen DNeasy blood and tissue kit, Qiagen Cat No./ID: 69504 following manufacturer’s instructions. The study protocol was approved through Institutional Animal Ethics Committee (approval no. CPCSEA/IAEC/P-6/2018) which was formed in accordance with the norms of the Committee for the Purpose of Control and Supervision of Experiments on Animals (CPCSEA), Government of India and complied with standard guidelines on handling of experimental animals.

According to manufacturer’s instructions, 20 μl Proteinase K was pipetted into a 1.5 ml or 2 ml microcentrifuge tube, to this 50–100 μl anticoagulated blood was added and the volume was adjusted to 220 μl with PBS. 200 μl Buffer AL was added and mixed thoroughly by vortexing, and incubated at 56°C for 10 min. Finally 200 μl ethanol (96–100%) was added to the sample, and mixed thoroughly by vortexing. The mixture was added to the DNeasy Mini spin column placed in a 2 ml collection tube and centrifuged at 6000 x g for 1 min. Further the DNeasy Mini spin column was placed in a new 2 ml collection tube (previous flow through and collection tube was discarded), 500 μl Buffer AW1 was added and centrifuged for 1 min at 6000 x g. Again flow-through and collection tube was discarded. The DNeasy Mini spin column was placed in a new 2 ml collection tube, 500 μl Buffer AW2 was added, and centrifuged for 3 min at 20,000 x g to dry the DNeasy membrane. Flow-through and collection tube was discarded. DNeasy Mini spin column was placed in a clean 1.5 ml or 2 ml microcentrifuge tube and 200 μl Buffer AE was pipetted directly onto the DNeasy membrane. Sample was incubated at room temperature for 1 min, and then centrifuged for 1 min at 6000 x g to elute.

### Human tissue samples

To build the human-rat clock, we analyzed previously generated methylation data from n=850 human tissue samples (adipose, blood, bone marrow, dermis, epidermis, heart, keratinocytes, fibroblasts, kidney, liver, lung, lymph node, muscle, pituitary, skin, spleen) from individuals whose ages ranged from 0 to 93. The tissue samples came from three sources. Tissue and organ samples from the National NeuroAIDS Tissue Consortium ^52^. Blood samples from the Cape Town Adolescent Antiretroviral Cohort study ^53^. Skin and other primary cells provided by Kenneth Raj ^41^. Ethics approval (IRB#15-001454, IRB#16-000471, IRB#18-000315, IRB#16-002028).

### Plasma fraction experimental procedure

Although transfusion technologies for humans are well-developed and safe, transfusion of small animals is still at the infancy stage of development, requiring state-of-the-art techniques and remains challenging. We used a unique plasma fraction “Elixir” developed by Nugenics Research. The necessary dose of plasma fraction treatment was prepared and injected intravenously using sterile saline as vehicle. The calculated doses were administered intravenously to the animals of old treated group; 4 injections every alternate day for 8 days, and a second dosing starting from the 95^th^ day consisting of 4 injections every alternate day for 8 days, as shown in Supplementary Figure 4. Similar amount of sterile saline solution (placebo) was administered to the animals of Old control group. Body weight, food and water intake of the animals were monitored at each time point. Cognitive abilities of animals were evaluated using Barnes Maze apparatus (spanning a week of training) 1, 2, 3 and 4 months from the start of the 1^st^ series of injections. Blood samples were withdrawn at predetermined time intervals by retro orbital plexus during the treatment for haematological evaluation. Serum was separated from the blood samples of each animal and evaluated for biochemical parameters. Plasma was separated from the blood samples of each animal and was used for evaluation of inflammatory markers i.e. TNF-α and IL-6. Animals were sacrificed from each group at 155^th^ day of treatment and vital organs (brain, heart, lung and liver) of these animals were harvested for testing of oxidative stress biomarkers, level of Nrf2, histopathological and immunohistochemistry studies.

### End-point evaluations

#### Body Weight

Body weights of rats were recorded before the initiation of treatment protocol and then 30, 60, 90,120 and 155^th^ day.

#### Grip strength

A grip strength meter was used to measure forelimb grip strength which represents the muscle strength of animals. Briefly, as rat grasped the bar of muscle strength meter, the peak pull force was recorded on a digital force transducer. Tension was recorded at the time the rat released its forepaws from the bar. Six consecutive measurements were taken per day at intervals of one-minute.

#### Barnes Maze Learning Ability

The Barnes maze platform (91 cm diameter, elevated 90 cm from the floor) consisted of 20 holes (each 5 cm in diameter). All holes were blocked except for one target hole that led to a recessed escape box. Spatial cues, bright light, and white noise were used to motivate the rat to find the escape during each session. For the adaptation phase, each rat explored the platform for 60 s. Any rat that did not find the escape box was guided to it and remained there for 90 s. For the acquisition phase, each trial followed the same protocol, with the goal to train each rat to find the target and enter the escape box within 180 s. Rat remained in the box for an additional 60 s. Four trials per day, approximately 15 min apart, was performed for 6 consecutive days. (Flores et al., 2018) A Barnes maze apparatus used to determine the learning ability of the animals upon treatment. Performed in each month of experiment.

#### Haematological tests

Blood was collected from the retro-orbital plexus using heparinized capillary tubes before the treatment and on 60^th^ and 155^th^ day of the experiment. One portion of the blood was kept in plain bottles from which serum was collected and stored for biochemical analysis. The other portion was directly subjected for the estimation of various haematological parameters using standard instruments. The levels of haemoglobin (Hb), red blood cell count (RBC), packed cell volume (PCV), mean corpuscular volume (MCV), mean corpuscular haemoglobin (MCH), mean corpuscular haemoglobin concentration (MCHC) and platelets were analyzed in the blood samples in all experimental groups.

#### Biochemical test

Blood samples were collected from the retro-orbital plexus using heparinized capillary tubes before the treatment and on 30^th^, 60^th^, 90^th^, 125^th^ and 155^th^ day of the experiment. One portion of the blood was kept in plain bottles from which serum was collected and stored for biochemical analysis. Further, the levels of serum glutamate pyruvate transaminase (S.G.P.T-IU/L) were carried out by kinetic method recommended by International Federation of Clinical Chemistry (IFCC). All the tests were performed with commercially available diagnostic kits (Erba Mannheim, Germany on Erba Mannheim biochemistry semi auto analyzer). Kidney function tests such as determination of serum creatinine (mg/dl) and uric acid (mg/dl) levels were done according to modified Jaffe’s reaction with commercially available diagnostic kits (Erba Mannheim, Germany on Erba Mannheim biochemistry semi auto analyzer). Blood glucose level (Random) (mg/dl) (Gaikwad et al.,2015), Total protein (g/dl), Total Bilirubin (mg/dl), Direct Bilirubin (mg/dl), Triglyceride (mg/dl), HDL (mg/dl), Cholesterol (mg/dl), Albumin (g/dl) (Erba Mannheim) were determined.

#### Oxidative stress evaluation

At the end of experiment, brain, heart, lung and liver were isolated and 10% tissue homogenate was prepared in ice-cold 50 mM PBS (pH 7.4) by using homogenizer followed by sonication for 5 min. The homogenate was centrifuged at 2000 g for 20 min at 4°C and the aliquots of the supernatant were collected and stored at −20° C up to further evaluation.

#### Estimation of extent of lipid peroxidation (LPO) (Malondialdehyde (MDA)

The brain, heart, lung and liver tissue homogenate samples were treated with 1 % phosphoric acid solution and aqueous solution of 0.6 % thiobarbituric acid. The reaction mixture was heated at 80⁰ C for 45 minutes, cooled in an ice bath and extracted with 4.0 ml of N-butanol. The n-butanol layer was separated and the absorbance of the pink complex formed was estimated at 532 nm as an indicator of extend of lipid peroxidation.

#### Estimation of reduced glutathione (GSH)

The GSH content in the brain, heart, lung and liver tissue homogenate was determined by treating the homogenate with sulfhydryl reagent 5,5’-dithio-bis(2-nitrobenzoic acid) (DTNB) method. Briefly, 20 μl of tissue homogenate was treated with 180 μl of 1mM DTNB solution at room temperature. The optical density of resulting yellow color was measured at 412 nm using a microplate spectrophotometer (Powerwave XS, Biotek, USA).

#### Determination of the catalase activity

The brain, heart, lung and liver tissue homogenate (20 μl) was added to 1 ml of 10mM H2O2 solution in the quartz cuvette. The reduction in optical density of this mixture was measured by using spectrophotometer in UV mode at 240nm. Rate of decrease in the optical density across three minutes from the addition of heart homogenate was taken as an indicator of the catalase activity present in the homogenate.

#### Estimation of superoxide dismutase (SOD) activity

The brain, heart, lung and liver tissue homogenate (20 μl) was added to a mixture of 20 μl of 500 mM of Na_2_CO_3_, 2 ml of 0.3 % Triton X-100, 20 μl of 1.0 mM of EDTA, 5 ml of 10 mM of hydroxylamine and 178 ml of distilled water. To this mixture, 20 μl of 240 μM of NBT was added. The optical density of this mixture was measured at 560 nm in kinetic mode for 3 minutes at one minute intervals. The rate increase in the optical density was determined as indicator of the SOD activity.

#### Nrf2 concentration in vital organs

Nrf2 was estimated in brain, heart, lung and liver homogenates using Nrf2 ELISA kit (Kinesis Dx, USA),. Organ was removed; and homogenate was prepared and kept at −20°C until the execution of assay. The Nrf2 level was determined by using kit according to the manufacturer’s protocol, and the values were calculated from the optical density of samples.

#### Pro-inflammatory cytokines (IL-6 and TNF-α)

The cytokines were estimated in plasma, which was separated from blood of animals and kept at −20°C until the execution of assay. The pro-inflammatory cytokine levels including TNF-α and IL-6 were determined by using sandwich ELISA kit (Kinesis Dx, USA), according to the manufacturer’s protocol, and the values were calculated from the optical density.

#### Histopathology of Vital Organs

Brain, heart, spleen, kidney, lung, liver and testis tissues fixed in neutral buffered 10% formalin solution were embedded in paraffin, and serial sections (3 μm thick) were cut using microtome (Leica RM 2125, Germany). The representative sections were stained with hematoxylin and eosin and examined under light microscope (Leica, Germany). The Histopathological data was objective and the sections were screened from a pathologist blinded to the treatments.

#### SA-β-gal staining

This assay was performed using a commercially available senescence β-Galactosidase staining detection kit (Cell Signaling, #9860). Briefly, cryosections were fixed with fixative solution for 10–15 min at room temperature, followed by staining with fresh β -gal staining solution overnight at 37°C. While the β –galactosidase is still on the plate, check the section under microscope (100X magnification) for the development of the blue color.

#### Oil Red O staining

Cryosections (6 μm thick) were fixed in neutral buffered 10% formalin solution for 10 min. The slides were incubated with freshly prepared Oil Red O working solution for 15 min. Lipid accumulation was digitalized using a microscope.

### DNA methylation profiling

We generated DNA methylation data using the custom Illumina chip “HorvathMammalMethylChip40”. By design, the mammalian methylation array facilitates epigenetic studies across mammalian species (including rats and humans) due to its very high coverage (over thousand X) of highly-conserved CpGs in mammals. Toward this end, bioinformatic sequence analysis was employed to identify 36 thousand highly conserved CpGs across 50 mammalian species. These 36k CpGs exhibit flanking sequences that are highly conserved across mammals. In addition, the custom array contains two thousand probes selected from human biomarker studies. Not all 36k probes on the array are expected to work for all species, but rather each probe is designed to cover a certain subset of species, such that overall all species have a high number of probes. The particular subset of species for each probe is provided in the chip manifest file which has been posted on Gene Expression Omnibus. The SeSaMe normalization method was used to define beta values for each probe^54^.

### Penalized Regression models

We developed the six different epigenetic clocks for rats by regressing chronological age on all CpGs that are known to map to the genome or *Rattus norvegicus*. Age was not transformed. We used all tissues for the pan-tissue clock. We restricted the analysis to blood, liver, and brain tissue for the blood, liver, and brain tissue clocks, respectively. Penalized regression models were created with the R function “glmnet” ^55^. We investigated models produced by both “elastic net” regression (alpha=0.5). The optimal penalty parameters in all cases were determined automatically by using a 10 fold internal cross-validation (cv.glmnet) on the training set. By definition, the alpha value for the elastic net regression was set to 0.5 (midpoint between Ridge and Lasso type regression) and was not optimized for model performance. We performed a cross-validation scheme for arriving at unbiased (or at least less biased) estimates of the accuracy of the different DNAm based age estimators. One type consisted of leaving out a single sample (LOOCV) from the regression, predicting an age for that sample, and iterating over all samples.

### Relative age estimation

To introduce biological meaning into age estimates of rats and humans that have very different lifespan; as well as to overcome the inevitable skewing due to unequal distribution of data points from rats and humans across age range, relative age estimation was made using the formula: Relative age= Age/maxLifespan where the maximum lifespan for rats and humans were set to 3.8 years and 122.5 years, respectively.

### Final version of epigenetic clocks

The final versions of our epigenetic clocks are meant for future studies of rat tissue samples. These final versions of clocks were developed by combining the original training data (n=517 rat tissues) with the “untreated” samples from the rat test data. Increasing the sample size of the training data leads to a higher accuracy according to a cross validation analysis (**Supplementary Figure 10, Supplementary Figure 11**). Using the final version of the epigenetic clocks, we find that the treatment effects become even more significant especially for the hypothalamus **(Supplementary Figure 12**). Final versions of the pan-tissue clock, liver clock, blood clock, brain clock, and “human-rat” clock can be found in Supplementary Material.

### Epigenome wide association studies (EWAS) of age

EWAS was performed in each tissue separately using the R function “standardScreeningNumericTrait” from the “WGCNA” R package ^56^. Next, the results were combined across tissues using Stouffer’s meta-analysis method. Our epigenome wide association test studies of chronological age reveal that aging effects in one tissue are often poorly conserved in another tissue (**Supplementary Figure 13**).

## URLs

## Acknowledgements

The development of the rat tissue clocks was supported by the Paul G. Allen Frontiers Group (SH) and a grant from Open Philanthropy (SH). The heterogeneous stock rats provided by (HC and LS) were supported by NIH grant DA-037844 (NIDA, HC and LW). RG was supported by grant # MRCF 7-25-19 from the Medical Research Charitable Foundation and the Society for Experimental Gerontological Research, New Zealand (RG). Human tissue sample collection was supported by NIH funding through the NIMH and NINDS Institutes by the following grants: Manhattan HIV Brain Bank (MHBB): U24MH100931; Texas NeuroAIDS Research Center (TNRC): U24MH100930; National Neurological AIDS Bank (NNAB): U24MH100929; California NeuroAIDS Tissue Network (CNTN): U24MH100928 Data Coordinating Center (DCC): U24MH100925. Human blood samples were supported by R21MH107327. The contents are solely the responsibility of the authors and do not necessarily represent the official view of the NNTC or NIH.

## Authors contributions

The plasma fraction treatment was developed by Harold Katcher (HK) in consultation with Akshay Sanghavi (AS). KR, SH and HK drafted the manuscript. All authors helped to edit the article. JZ and SH developed the epigenetic clocks. SH generated the DNA methylation data. JZ, KR, SK, CL, SH carried out statistical analyses and created figures and tables. KS managed the plasma treatment project and carried out the experiments with SK and AgS. RG, HC, CH, LSW, MC-M, ML, PC, TW, AM contributed rat tissue samples. AL and SH contributed human data.

