## Supplementary Figures and Tables for "Reversing age: dual species measurement of epigenetic age with a single clock"

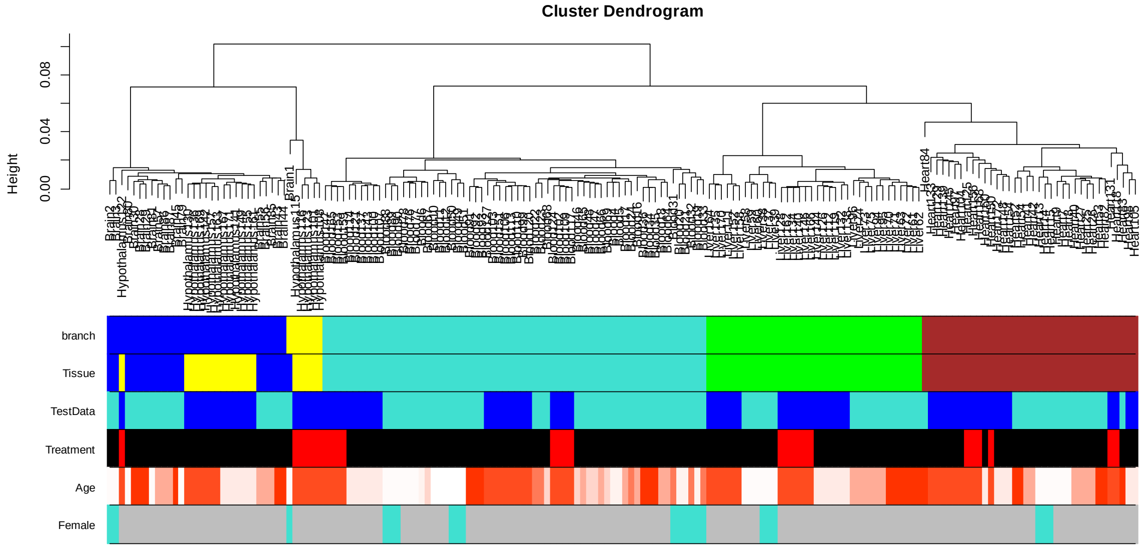


**Supplementary Figure 1. Unsupervised hierarchical clustering of tissue samples.** Branches largely correspond to tissue types. Although the data were generated from 3 different batches, the samples clustered by tissue type, showing negligible batch effects.

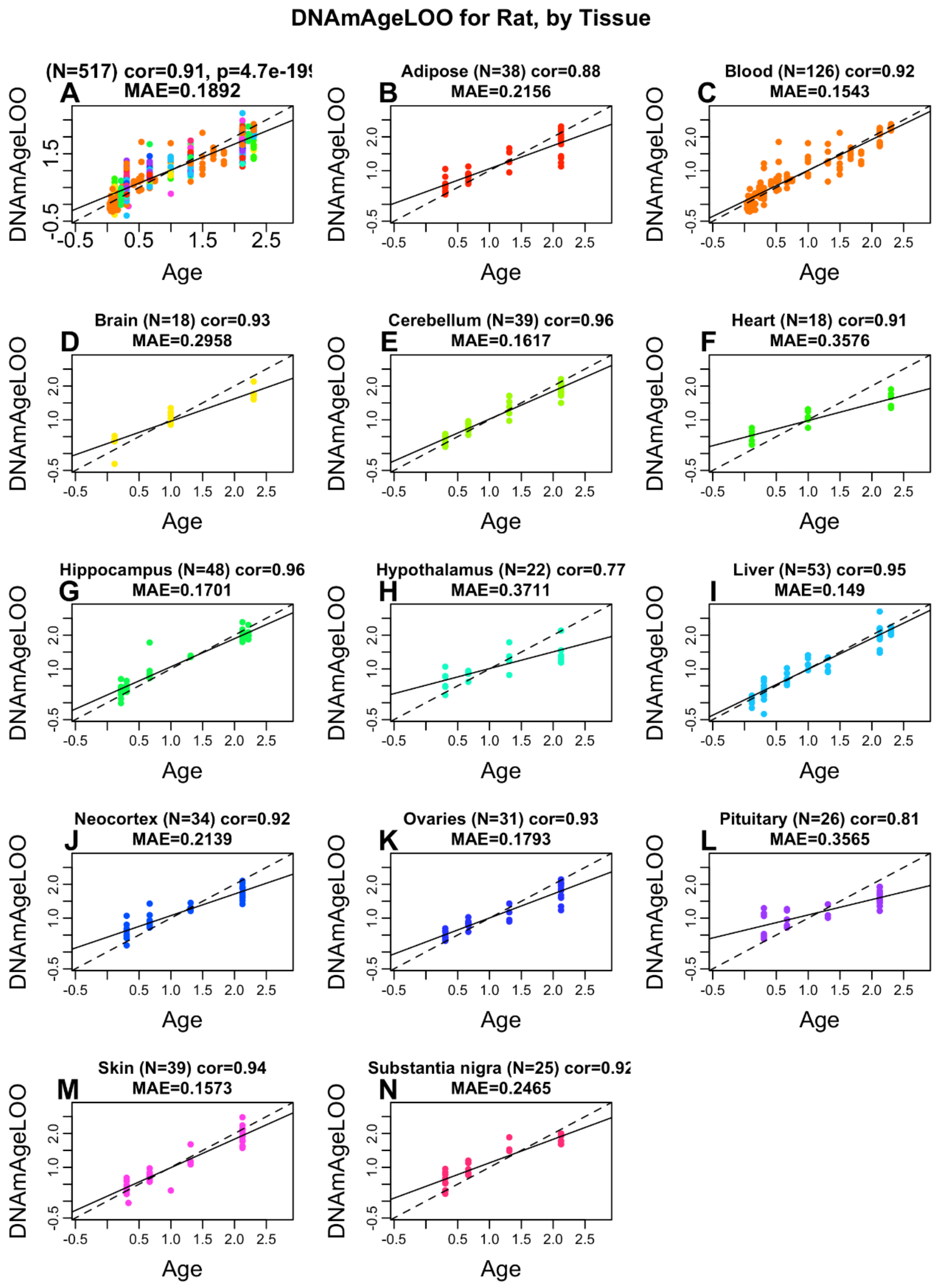


**Supplementary Figure 2. Pan tissue clock for rats applied to different tissues**. A) All tissues. B) adipose, C) blood, D) whole brain, E) cerebellum, F) heart, G) hippocampus, H) hypothalamus, I) liver, J) brain neocortex, K) ovaries, L) pituitary, M) skin, N) substantia nigra. The title of each panel reports the tissue, sample size, Pearson correlation coefficient and median absolute deviation (median error). Chronological age (x-axis) versus leave one sample out estimate of age.

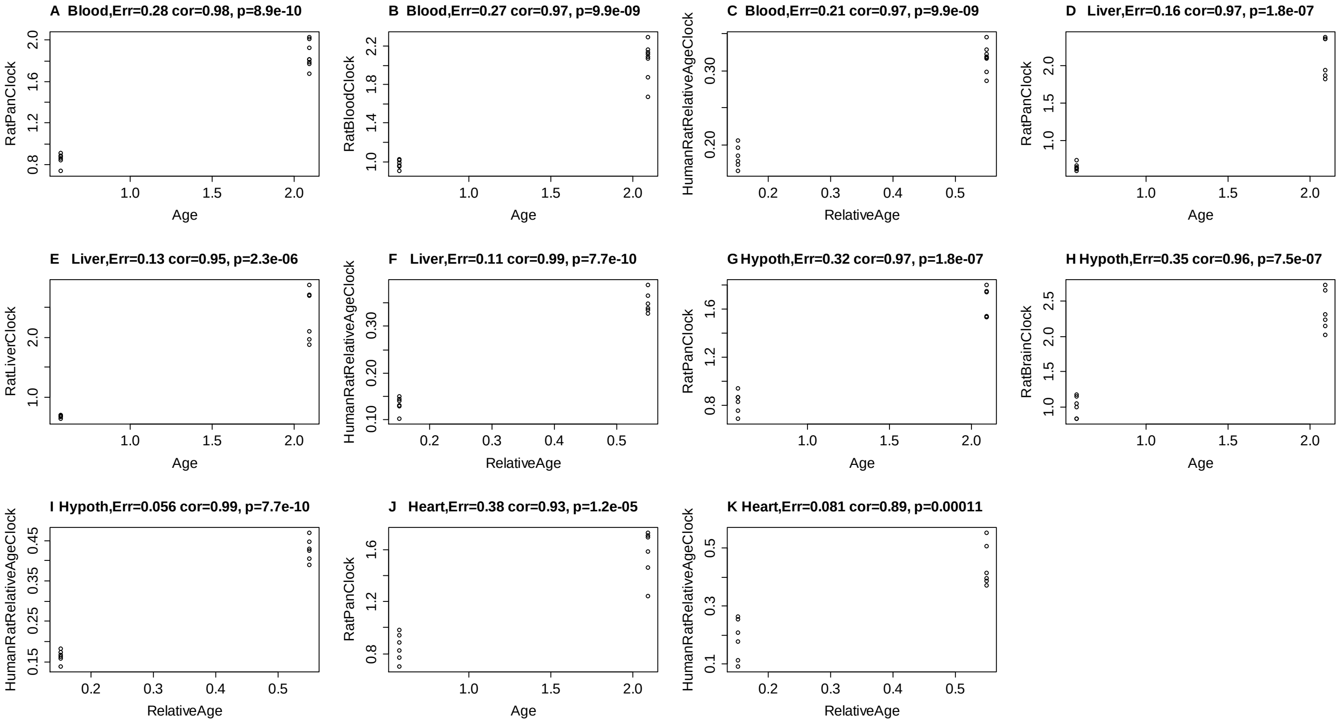

**Supplementary Figure 3: Epigenetic clocks applied to independent test data.** The six epigenetic clocks were applied to DNA profiles from un-treated samples from the plasma fraction study. The y-axis reports epigenetic age (in years) as measured by the indicated clocks. The heading reports the tissue type, the correlation between epigenetic and chronological age and the median error (in years). While the age correlations are high, the median errors are sub-optimal, up to 0.38 years. As such, the final versions of the clocks incorporated un-treated samples from the test data as well (Methods).

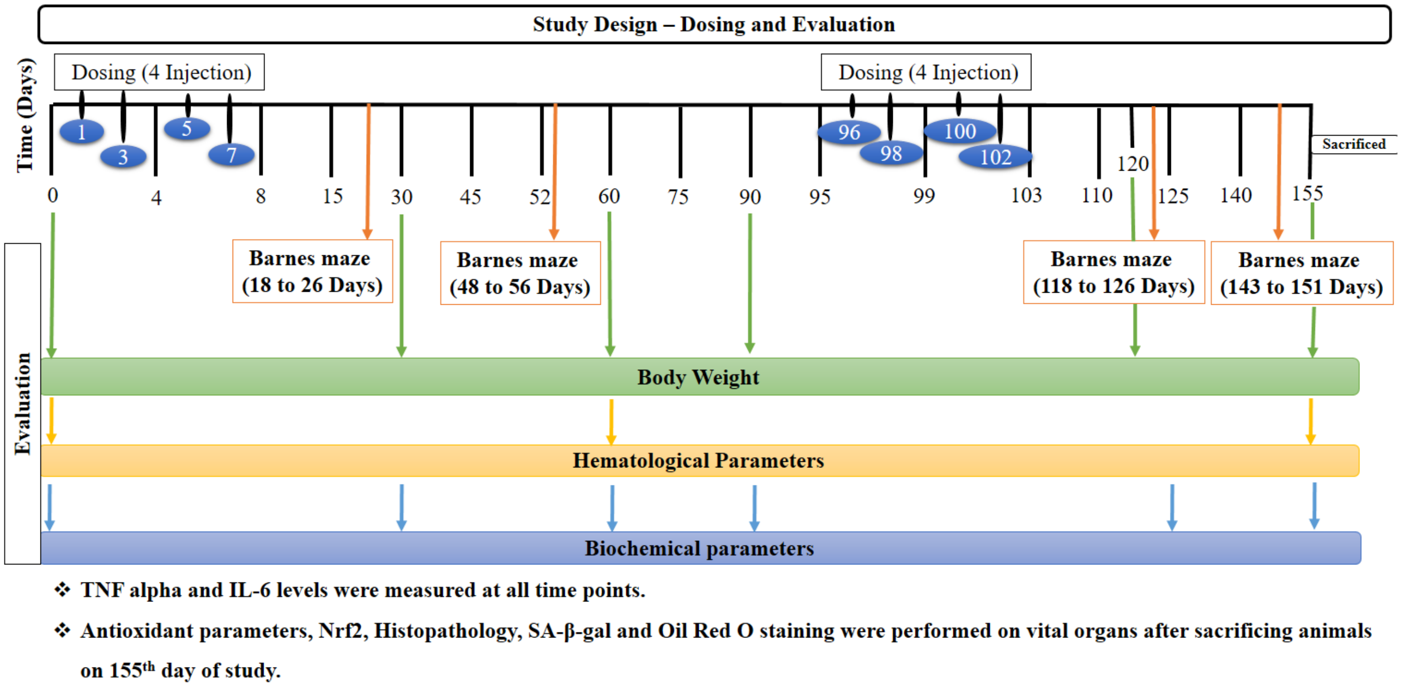


**Supplementary Figure 4:** Schematic representation of the study design with timeline indicating plasma fraction treatments as well as cognitive, physical, haematological and biochemical tests.

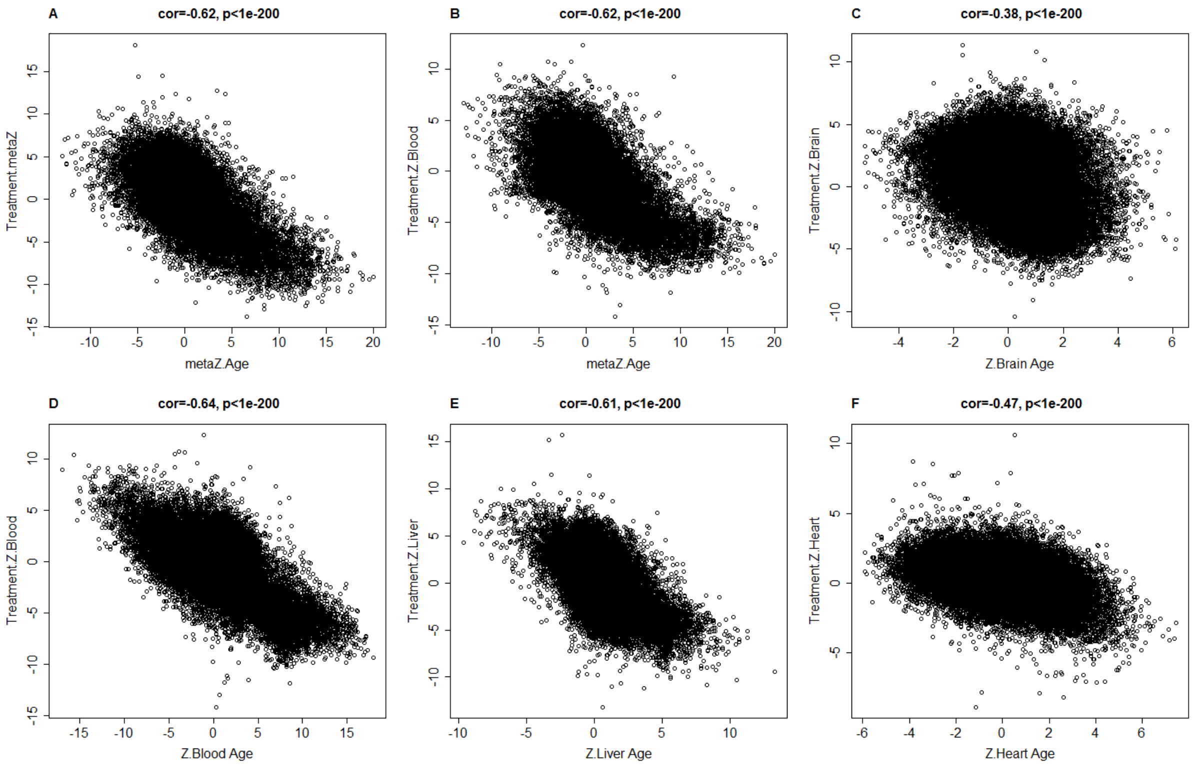


**Supplementary Figure 5: EWAS results for age versus EWAS results of plasma fraction treatment.** Each dot corresponds to a CpG on the mammalian array. The x axis reports Z statistics from a correlation test of CpG methylation versus chronological age. Positive/negative values correspond to positive/negative correlation coefficients with age, respectively. The y-axis reports a Z statistic for the treatment effect. Positive/negative values correspond to gain/loss of methylation associated with the treatment. A-B) The x axis corresponds to a meta-analysis of age effects across all rat tissues (Adipose, Blood, Brain, Cerebellum, Heart, Hippocampus, Hypothalamus, Liver, Neocortex, Ovaries, Pituitary, Skin, Substantia nigra). Age effects in C) Brain, D) Blood, E) Liver, F) Heart. A) Treatment effects across four tissues. Stouffer's meta-analysis Z statistic across hypothalamus, blood, liver, heart. Treatment effects in individual tissues B) Blood, C) Hypothalamus, D) Blood, E) Liver, F) Heart.

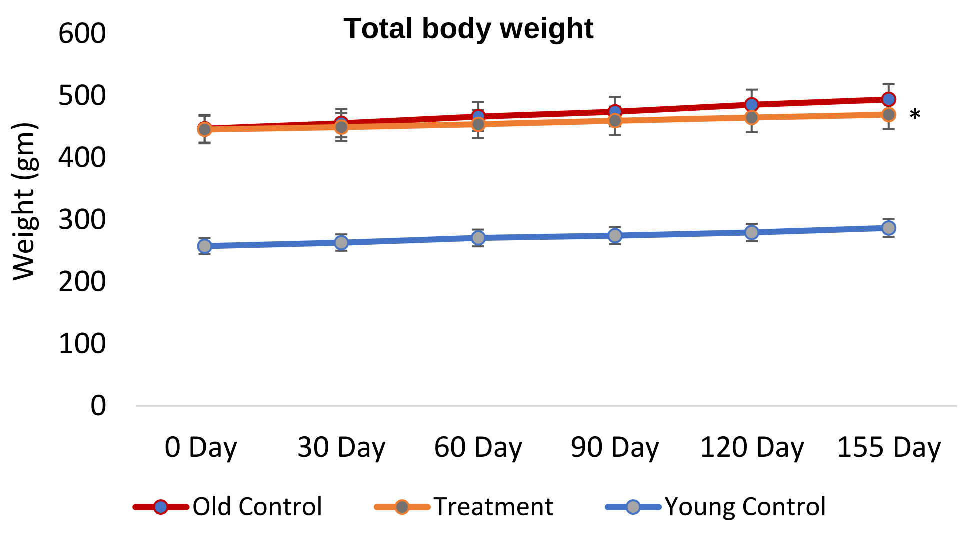


**Supplementary Figure 6A.** Weight of rats measured at regular intervals at indicated times during the 155-day experiment. Each group measurement was from 6 rats.

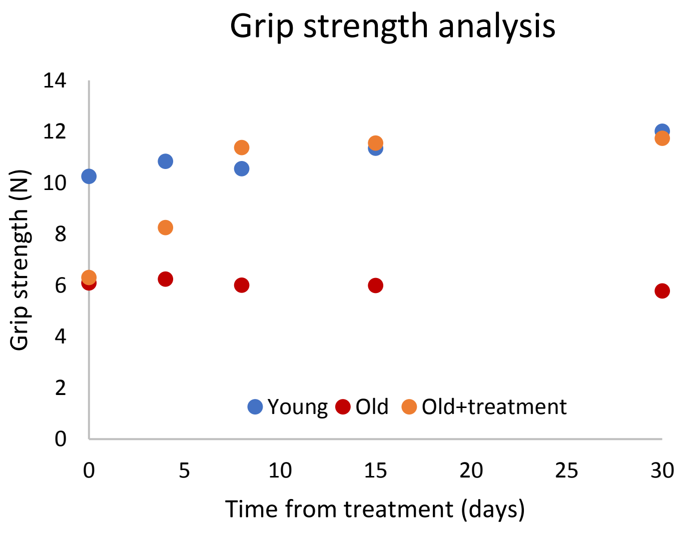


**Supplementary Figure 6B.** Measurement of grip strength of the indicated rat groups at various times post-treatment. Each group consists of 6 rats. For clarity, the plotted data points represent average values from 6 rats each. Detailed measurements of each parameter with standard deviations are provided in Supplementary table 5.

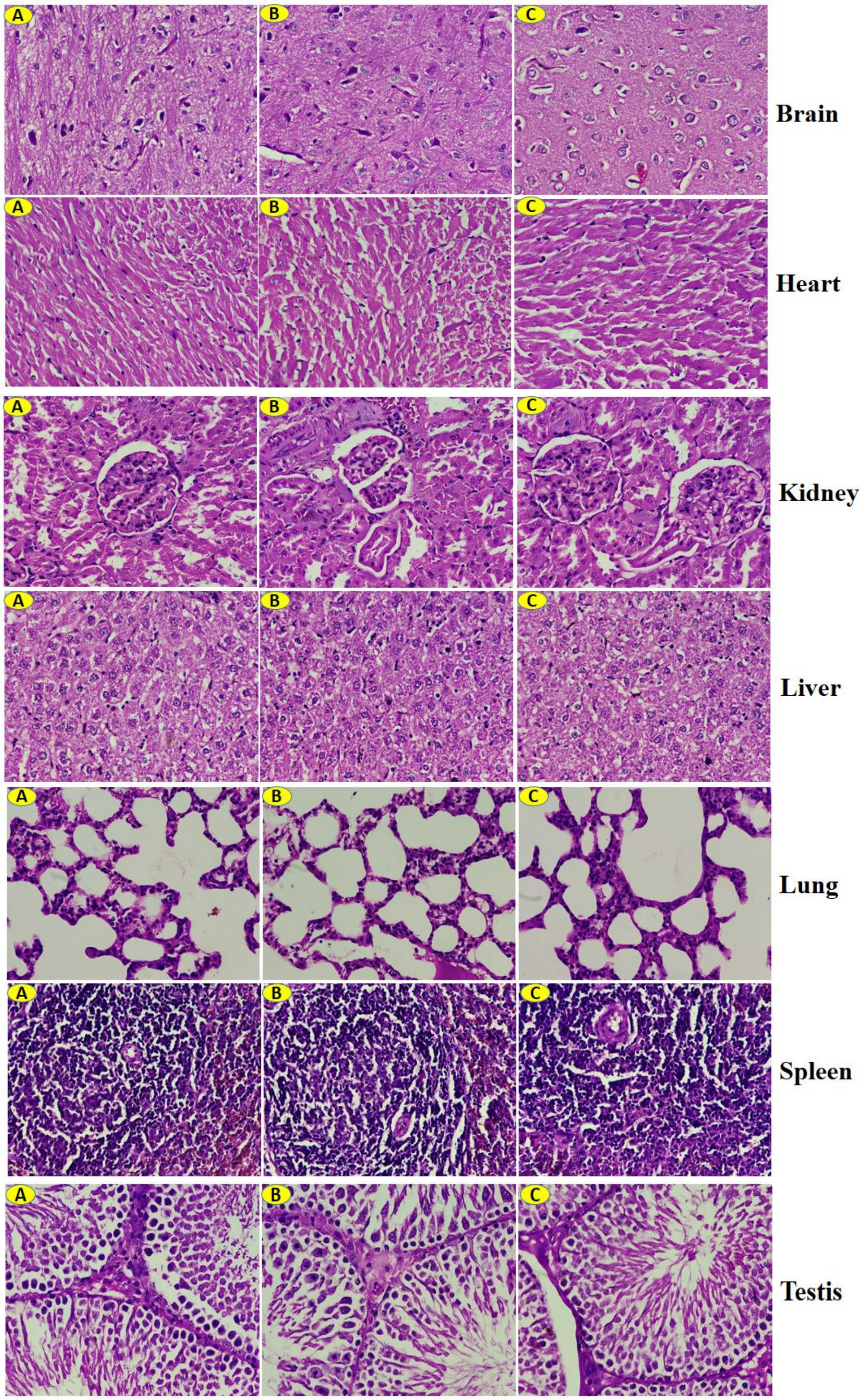


**Supplementary Figure 7:** Histological analyses of vital organs and tissues of rats employed in experiment. Images of the left column (A) are from old rats; the right column (C) are images of tissues from young rats while the middle column (B) are images of tissues from old rats treated with plasma fraction. Results of histopathological examination are tabulated in Supplementary Table 4

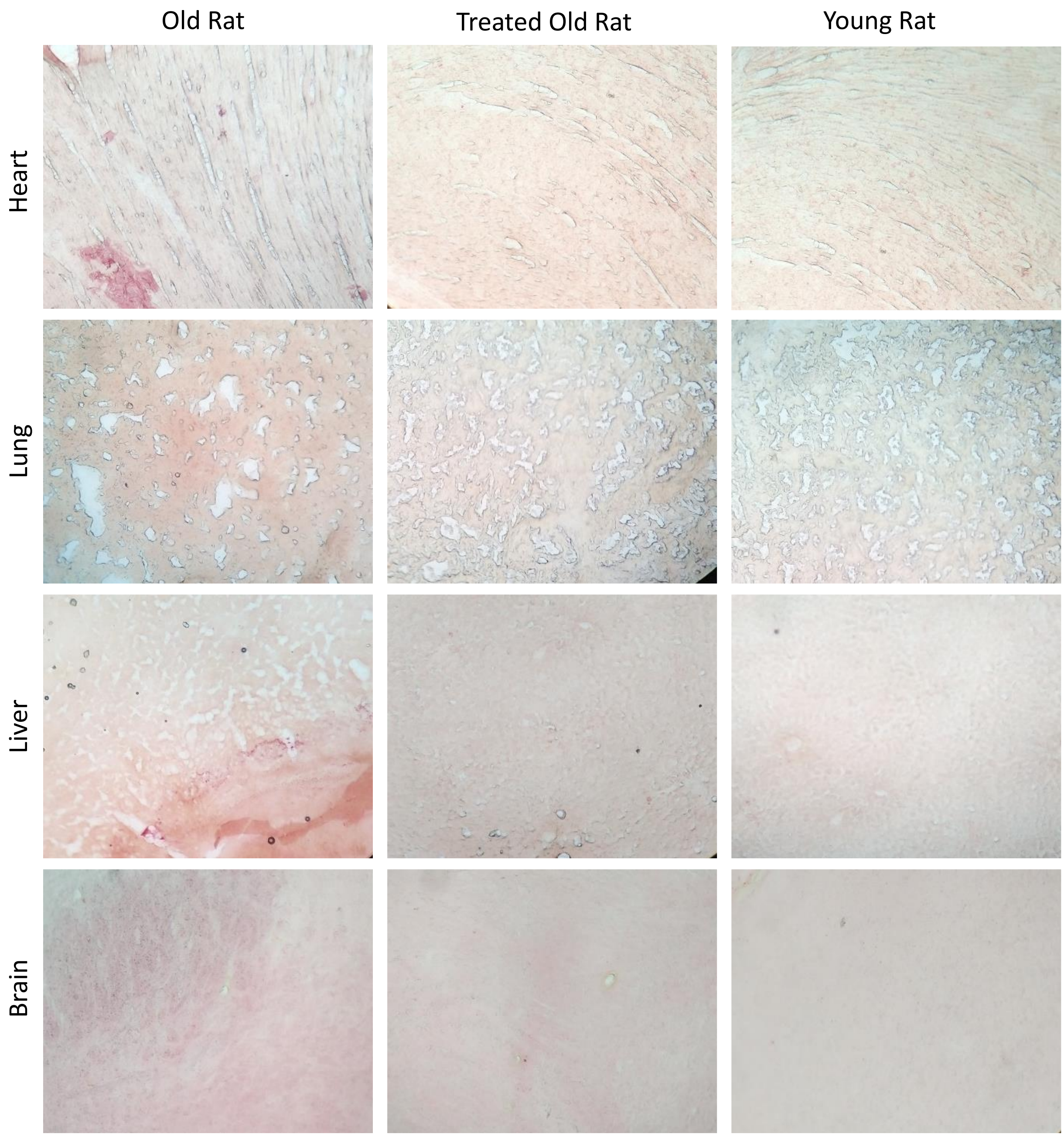


**Supplementary Figure 8:** Tissues from old untreated rats, Plasma fraction-treated old rats and young rats were subjected to Oil Red O staining to reveal accumulation of fat in tissues.

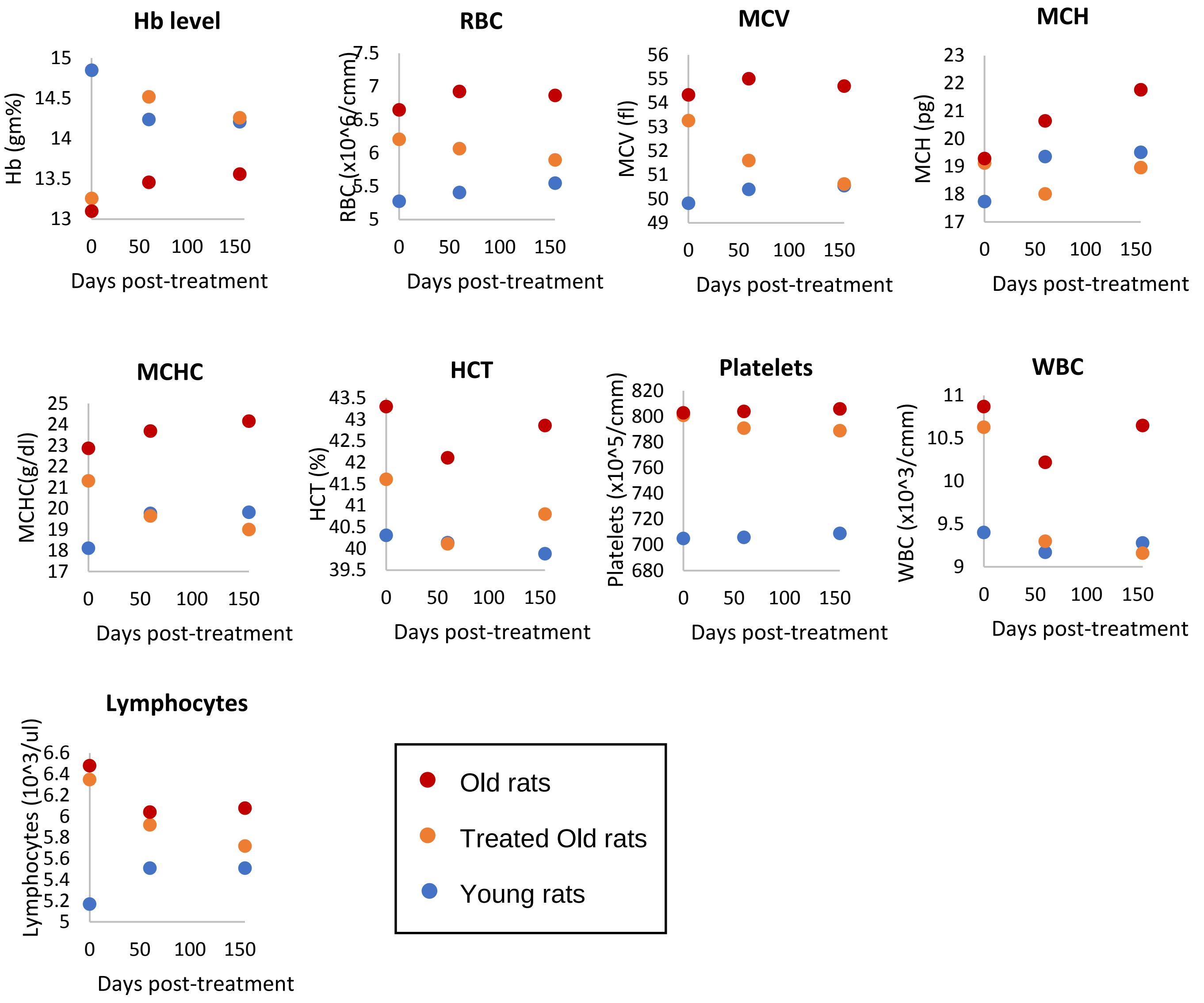


**Supplementary Figure 9:** Effects of plasma fraction treatment on haematological properties of rats at 0, 60 and 155 days from the start of experiment. Hb = haemoglobin, RBC = red blood cell count, MCV = mean corpuscular volume, MCH = mean corpuscular haemoglobin, MCHC = mean corpuscular haemoglobin concentration, HCT = haematocrit and WBC = white blood cells. Red dots represent data points of old rats, orange dots represent treated old rats and blue represent young rats. For clarity, the plotted data points represent average values from 6 rats each. Detailed measurements of each parameter with standard deviations are provided in Supplementary table 6.

**Leave-One-Out Analysis of All Final Epigenetic Clocks**

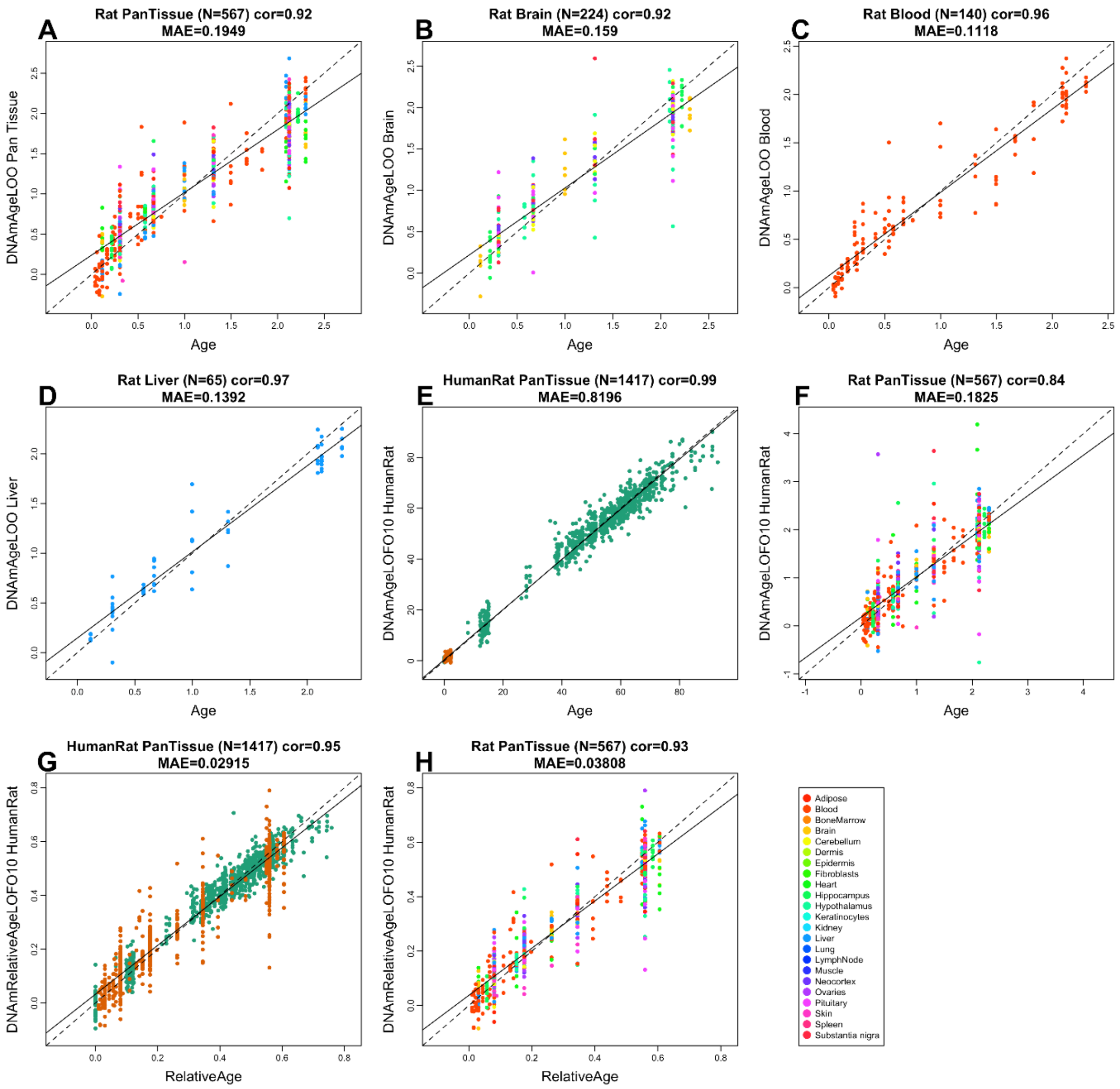


**Supplementary Figure 10:** This figure is analogous to Figure 1, but it reports estimates of the predictive accuracy of the final version of the rat clocks. Cross-validation was carried out on an increased number of rat tissues (n=567) by combining the original training data (n=517) with rat tissues from untreated animals of the test data set.

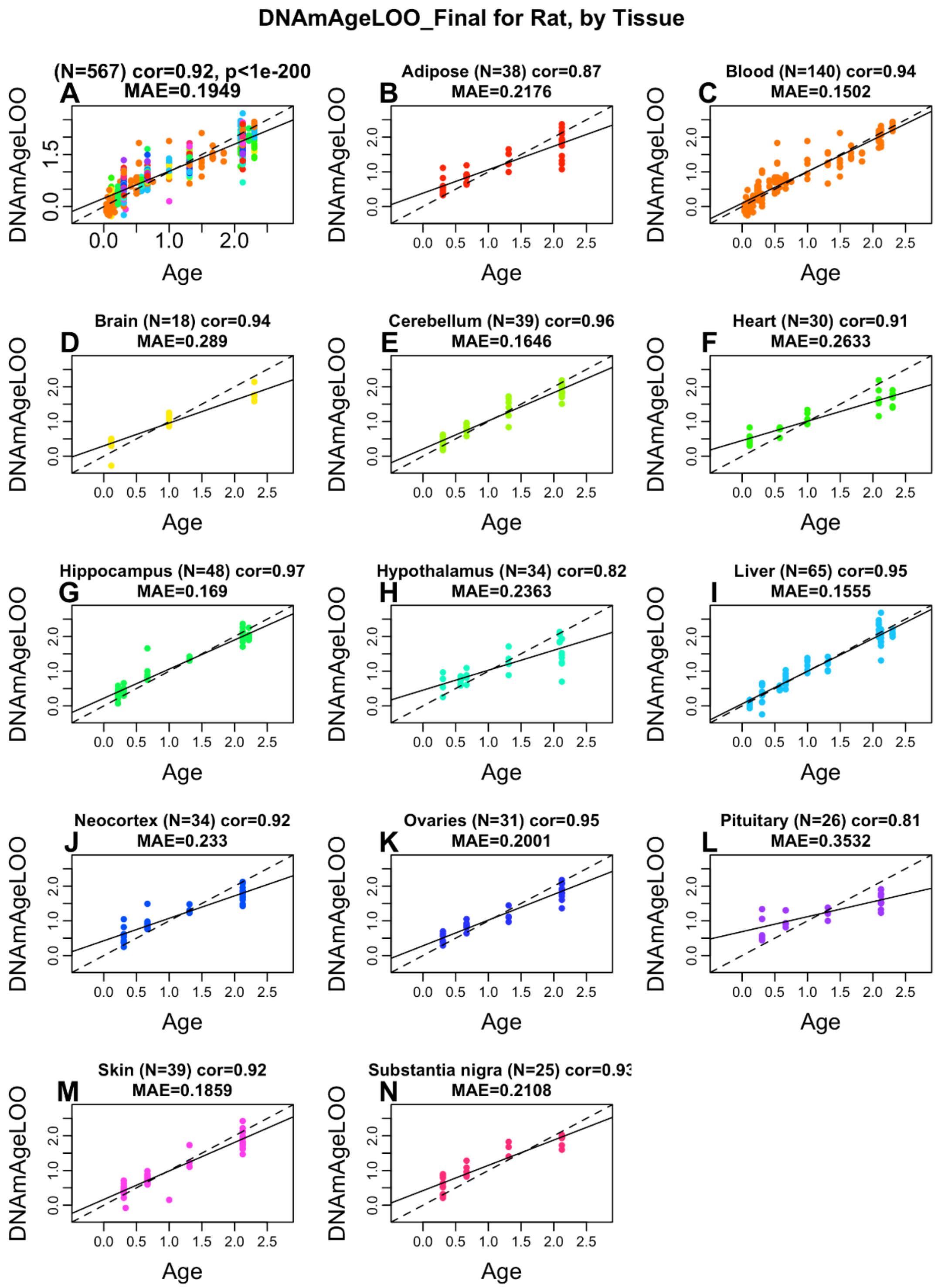


**Supplementary Figure 11**: Estimating the accuracy of the final version of the pan tissue rat clock. Each panel corresponds to a different source of DNA in rat tissue. Leave one out cross validation estimates of DNAmAge (y-axis) versus chronological age.

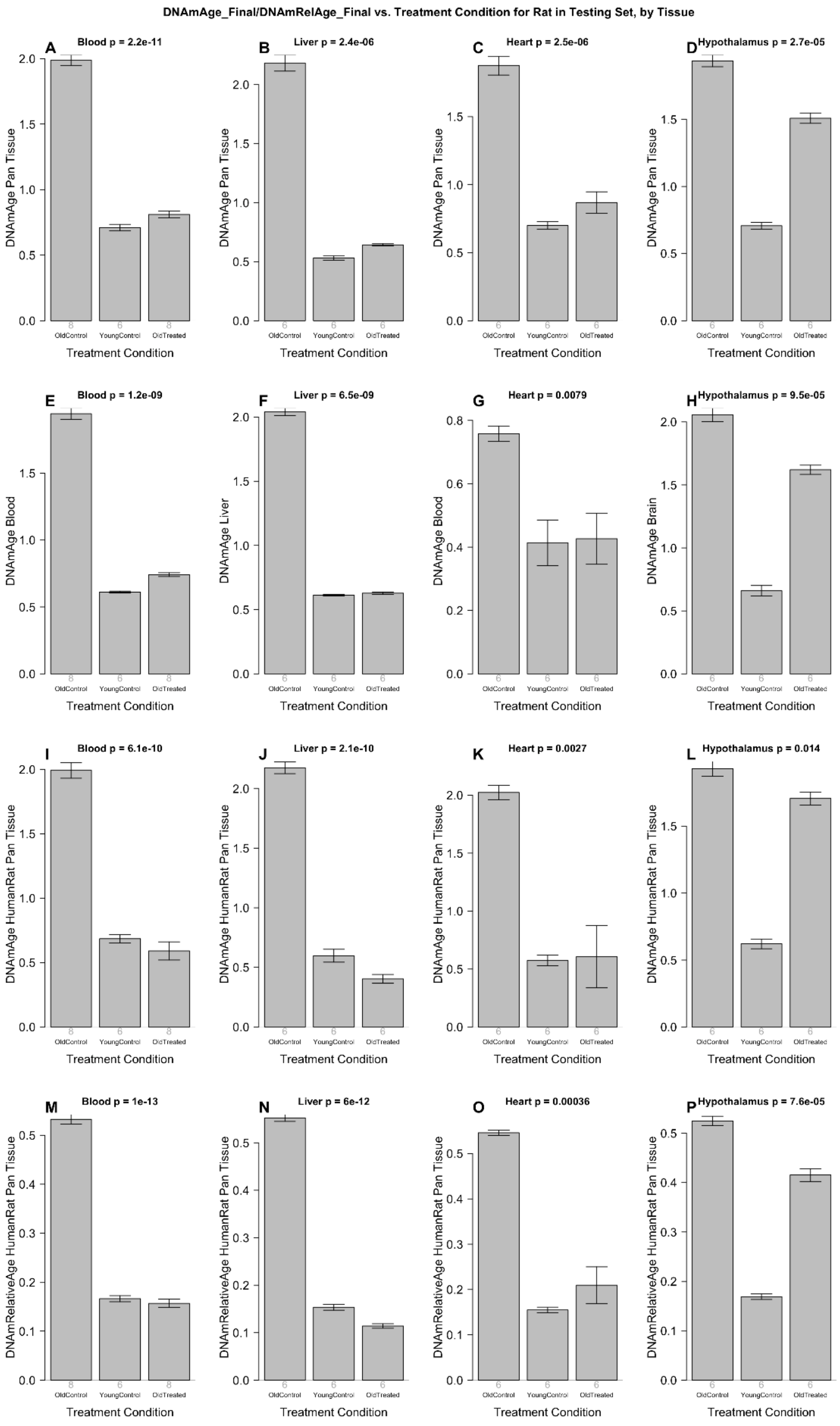


**Supplementary Figure 12**. **Epigenetic clock analysis of plasma fraction treatment based on the final version of epigenetic clocks.**This figure is analogous to Figure 2, but it reports results for the final version of the epigenetic clock based on an increased number of rat tissues (n=567) (resulting from combining the original n=517 original training data with rat tissues from untreated animals of the test data set). Six epigenetic clocks applied to independent test data from four rat tissue type (columns): blood, liver, heart, and hypothalamus. A-D) Rat pan-tissue clock. E) Rat blood clock applied to blood. F) Rat liver clock applied to liver. G) Rat blood clock applied to heart. H) Rat brain clock applied to hypothalamus. I-L) Human-rat clock measure of absolute age. M-P) Human-rat clock measure of relative age defined as age/maximum species lifespan. Each bar-plot reports the mean value and one standard error. Student T-test p values result from a 2 group comparison of old controls (left bar) versus old treated samples (right bar), i.e. the young controls were omitted.

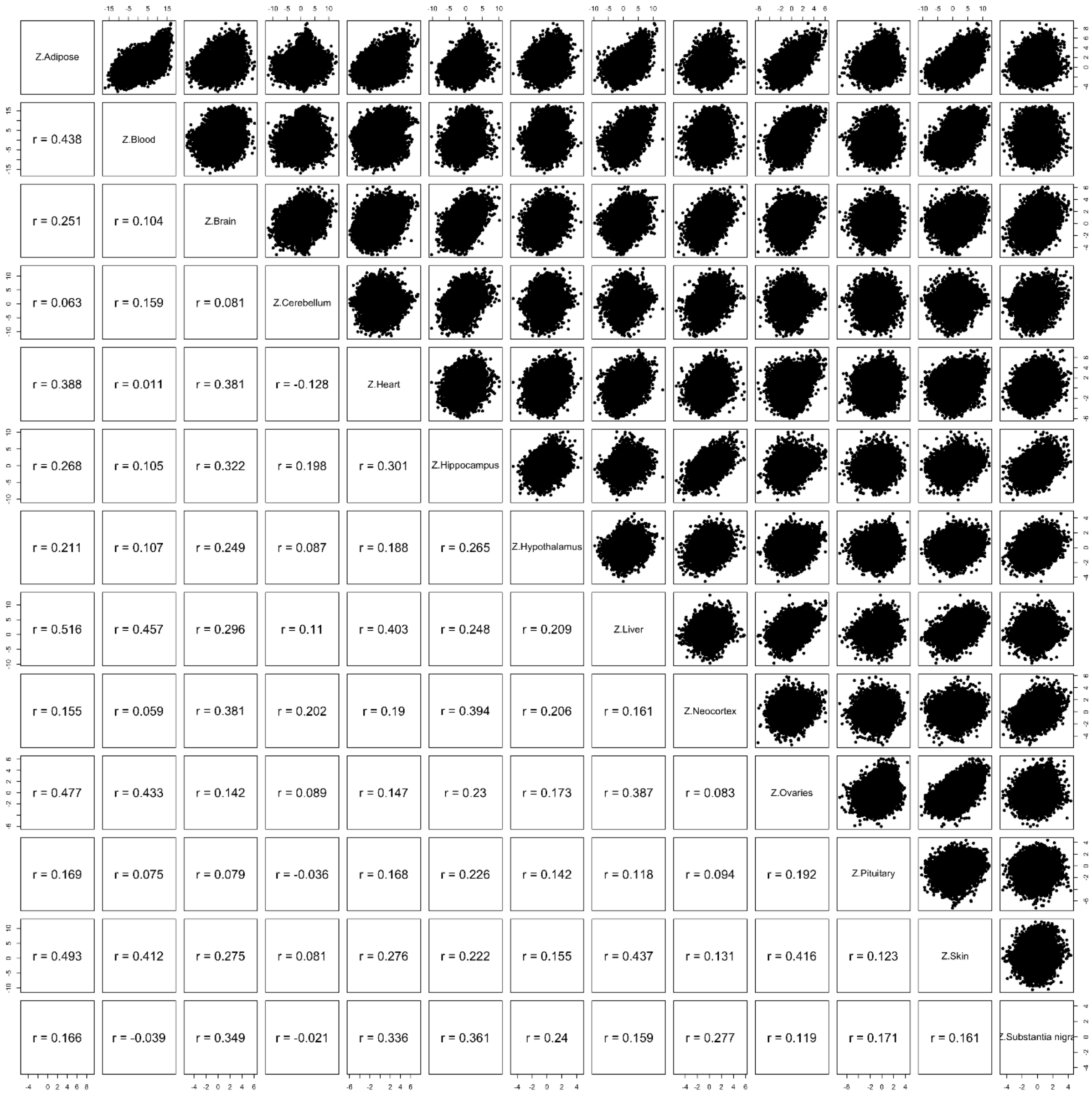


**Supplementary Figure 13**. EWAS was performed in each rat tissue separately using the R function "standardScreeningNumericTrait" from the "WGCNA" R package. Z statistics from correlation tests in different tissues. Upper panel report scatter plots for Z statistics in different tissues. Lower panels: corresponding Pearson correlation coefficients.

**Supplementary Tables**

| Tissue | N | No. Female | Mean Age | Min. Age | Max. Age |
|---|---|---|---|---|---|
| Adipose | 38 | 38 | 1.15 | 0.305 | 2.12 |
| Blood | 148 | 72 | 0.931 | 0.0384 | 2.3 |
| Whole Brain | 18 | 3 | 1.29 | 0.115 | 2.3 |
| Cerebellum | 39 | 38 | 1.23 | 0.305 | 2.12 |
| Heart | 36 | 3 | 1.29 | 0.115 | 2.3 |
| Hippocampus | 48 | 48 | 1.13 | 0.216 | 2.22 |
| Hypothalamus | 40 | 22 | 1.22 | 0.305 | 2.12 |
| Liver | 71 | 37 | 1.22 | 0.115 | 2.3 |
| Neocortex | 34 | 34 | 1.16 | 0.305 | 2.12 |
| Ovaries | 31 | 31 | 1.22 | 0.305 | 2.12 |
| Pituitary | 26 | 26 | 1.26 | 0.305 | 2.12 |
| Skin | 39 | 37 | 1.16 | 0.305 | 2.12 |
| Substantia nigra | 25 | 25 | 1.08 | 0.305 | 2.12 |

**Supplementary Table 1.** Description of biological materials from which DNA methylation profiles were derived. The table reports the results for training (N=517) and test set (N=76) combined. Other supplementary tables provide details on the training and test data. N=Total number of tissues. Number of females. Age: mean, minimum and maximum in units of years.

| Sr. No | Age in Weeks | Age in Months | Number of Animal gender-wise (n=6) | Blood sample | Tissue samples | Total Samples |
|---|---|---|---|---|---|---|
| 1 | 3 | 0.7 | 3- Male and 3- Female | 6 | - | 6 |
| 2 | 6 | 1.4 | 3- Male and 3- Female | 6 | Heart, Liver, Brain (18 samples) | 24 |
| 3 | 12 | 2.8 | 3- Male and 3- Female | 6 | - | 6 |
| 4 | 28 | 6.4 | 6- Male | 6 | - | 6 |
| 5 | 52 | 12.0 | 6- Male | 6 | Heart, Liver, Brain (18 samples) | 24 |
| 6 | 78 | 18.0 | 3- Male and 3- Female | 6 | | 6 |
| 7 | 120 | 27.6 | 6- Male | 6 | Heart, Liver, Brain (18 samples) | 24 |
| **Total samples (42+54=96)** | | | | **42** | **54** | **96** |

**Supplementary Table 2.** Description of the rat tissue samples provided by Dr Kavita Singh for training of rat clock.

| Sr. No | Group | Age in Weeks | Age in Months | Number of Animal (n=6) | Blood sample | Tissue samples | Total Samples |
|---|---|---|---|---|---|---|---|
| 1 | Old Control | 109 | 27.0 | 6 Male | 6+2 replicates | Heart, Liver, Hypothalamus (6 samples each) | 26 |
| 2 | Old Treatment | 109 | 24.0 | 6 Male | 6+2 replicates | Heart, Liver, Hypothalamus (6 samples each) | 26 |
| 3 | Young Control | 30 | 4.0 | 6 Male | 6 | Heart, Liver, Hypothalamus (6 samples each) | 24 |
| | **Total samples (22+54=76)** | | | | **22** | **54** | **76** |

**Supplementary Table 3.** Description of animals employed for testing the effects of plasma fraction treatment

**Histopathological evaluation of rat tissues**

| Rat group | Liver | Lungs | Kidney | Heart | Spleen | Brian | Testes |
|---|---|---|---|---|---|---|---|
| **Old Control** | Granular degeneration of mild severity, Minimally multifocal minimal periportal mononuclear cell infiltration | NAD | NAD | NAD | NAD | NAD | NAD |
| **Old Control** | Granular degeneration of mild severity | NAD | NAD | NAD | NAD | NAD | NAD |
| **Old Control** | Granular degeneration of mild severity | NAD | NAD | NAD | NAD | NAD | NAD |
| **Treatment** | Granular degeneration of mild severity | NAD | NAD | NAD | NAD | NAD | NAD |
| **Treatment** | Granular degeneration of mild severity | NAD | NAD | NAD | NAD | NAD | NAD |
| **Treatment** | Granular degeneration of mild severity | NAD | NAD | NAD | NAD | NAD | NAD |
| **Young Control** | Granular degeneration of mild severity | NAD | NAD | NAD | NAD | NAD | NAD |
| **Young Control** | Granular degeneration of mild severity | NAD | NAD | NAD | NAD | NAD | NAD |
| **Young Control** | Granular degeneration of mild severity | NAD | NAD | NAD | NAD | NAD | NAD |

**Supplementary Table 4:** Results of histopathological analyses of rat tissues. Lesions suggestive of any toxicity were not noted. NAD= No Abnormalities Detected. Representative images of the tissues are shown in Supplementary figure 6.

| Groups | Grip Strength in N | | |
|---|---|---|---|
| | **Old control** | **Treatment** | **Adult Control** |
| **0 Day** | 6.10±0.32 | 6.31±0.44 | 10.25±1.01 |
| **4 Day** | 6.24±0.44 | 8.25±0.94 | 10.84±1.12 |
| **8 Day** | 6.01±0.46 | 11.38±0.83 | 10.55±0.99 |
| **15 Day** | 6.00±0.89 | 11.55±0.88 | 11.35±1.15 |
| **30 Day** | 5.78±0.75 | 11.74±0.76 | 12.01±1.23 |

**Supplementary Table 5:** Measurement (with standard deviations) of grip strength of indicated groups of 6 rats each. These average values formed the graphs in Supplementary figure 5B

| | **Groups** | Hb (gm %) | RBC (x $10^6$ /cmm) | WBC (X $10^3$ /cmm) | Platelets (X $10^5$/ cmm) | HCT (%) | MCV (fl) | MCH (pg) | MCHC g/dl | Lymphocytes ($10^3$ cells/µl) |
|---|---|---|---|---|---|---|---|---|---|---|
| 0 Day | Old | 13.10±0.8 | 6.65±0.5 | 10.87±1.0 | 803.00±6.2 | 43.03±1.0 | 54.34±0.9 | 19.29±0.8 | 22.86±0.7 | 6.48±0.8 |
| | Old treated | 13.26±1.0 | 6.21±0.6 | 10.63±1.2 | 801.83±5.3 | 41.61±2.1 | 53.27±1.5 | 19.13±0.7 | 21.31±1.2 | 6.35±1.1 |
| | Young | 14.85±0.9 | 5.28±0.9 | 9.40±0.7 | 705.33±8.0 | 40.31±0.6 | 49.83±1.3 | 17.74±1.0 | 18.10±1.1 | 5.17±0.7 |
| 60 Day | Old | 13.46±0.9 | 6.93±0.4 | 10.22±0.4 | 804.17±4.0 | 42.11±1.7 | 55.01±0.9 | 20.65±0.8 | 23.68±1.3 | 6.04±0.3 |
| | Old treated | 14.52±1.5 | 6.07±0.4 | 9.30±0.6 | 791.67±7.2 | 40.11±1.3 | 51.60±1.2 | 18.02±1.0 | 19.64±1.3 | 5.92±0.7 |
| | Young | 14.24±0.5 | 5.41±0.4 | 9.17±0.5 | 706.17±7.6 | 40.14±1.1 | 50.40±0.8 | 19.36±0.7 | 19.76±1.3 | 5.51±0.9 |
| 155 Day | Old | 13.56±0.5 | 6.87±0.6 | 10.65±0.5 | 806.50±5.1 | 42.86±0.8 | 54.71±0.9 | 21.77±1.0 | 24.16±0.8 | 6.08±0.2 |
| | Old treated | 14.26±0.9 | 5.90±06 | 9.16±0.3 | 789.00±3.5 | 40.80±1.4 | 50.63±1.3 | 18.97±0.7 | 18.99±0.8 | 5.72±0.5 |
| | Young | 14.21±0.7 | 5.55±0.6 | 9.28±0.9 | 709.83±7.7 | 39.88±1.0 | 50.55±0.7 | 19.52±0.7 | 19.81±1.0 | 5.51±0.7 |

**Supplementary Table 6:** Blood indices measurements (with standard deviation) at indicated time points post-treatment. Each measurement was taken from 6 rats per group. These average values formed the graphs in Supplementary figure 8.

**Detailed parameters of figure 3**

| Sr. No. | Parameter | Group | Old control | Treatment | Young Control |
|---|---|---|---|---|---|
| 1 | **Total Bilirubin (mg/dL)** | 0 Day | 0.90±0.10 | 0.90±0.11 | 0.56±0.08 |
| | | 30 Day | 0.92±0.09 | 0.83±0.12 | 0.55±0.09 |
| | | 60 Day | 0.92±0.11 | 0.82±0.10 | 0.57±0.08 |
| | | 90 Day | 0.97±0.11 | 0.78±0.09 | 0.57±0.08 |
| | | 125 Day | 1.02±0.12 | 0.72±0.09 | 0.59±0.08 |
| | | 155 Day | **1.08±0.12** | **0.68±0.07 ### *** | **0.60±0.07** |
| 2 | **Direct Bilirubin (mg/dL)** | 0 Day | 0.575±0.06 | 0.583±0.07 | 0.257±0.05 |
| | | 30 Day | 0.588±0.06 | 0.563±0.06 | 0.258±0.05 |
| | | 60 Day | 0.605±0.07 | 0.520±0.05 | 0.273±0.04 |
| | | 90 Day | 0.618±0.06 | 0.490±0.06 | 0.283±0.04 |
| | | 125 Day | 0.640±0.06 | 0.468±0.07 | 0.293±0.05 |
| | | 155 Day | **0.680±0.08** | **0.438±0.04 ### ***** | **0.308±0.04** |
| 3 | **Glucose (mg/dL)** | 0 Day | 173.0±5.57 | 174.9±4.97 | 152.0±8.80 |
| | | 30 Day | 174.4±5.13 | 169.6±2.88 | 152.3±8.34 |
| | | 60 Day | 178.7±5.02 | 167.5±3.53 | 155.4±8.69 |
| | | 90 Day | 180.0±5.37 | 165.3±3.00 | 157.9±9.60 |
| | | 125 Day | 183.2±5.04 | 163.5±3.59 | 161.2±9.55 |
| | | 155 Day | **186.0±3.32** | **163.6±4.10 ### *** | **164.6±10.09** |
| 4 | **Triglyceride (mg/dL)** | 0 Day | 57.2±10.16 | 56.4±10.45 | 25.0±9.01 |
| | | 30 Day | 70.5±5.45 | 45.1±6.67 | 28.9±8.80 |
| | | 60 Day | 79.3±5.52 | 45.4±5.84 | 30.8±7.83 |
| | | 90 Day | 84.1±5.63 | 43.6±6.05 | 33.1±7.25 |
| | | 125 Day | 91.4±5.08 | 41.7±5.78 | 34.7±7.92 |
| | | 155 Day | **103.7±5.30** | **38.0±5.64 ###** | **37.9±8.26** |
| 5 | **HDL (mg/dL)** | 0 Day | 109.7±9.19 | 111.3±8.95 | 142.5±6.71 |
| | | 30 Day | 110.7±7.05 | 117.2±8.44 | 144.3±5.92 |
| | | 60 Day | 108.9±6.89 | 127.0±8.16 | 145.1±6.25 |
| | | 90 Day | 108.0±7.67 | 130.6±9.13 | 147.2±6.70 |
| | | 125 Day | 105.9±7.78 | 138.2±10.11 | 149.0±6.49 |
| | | 155 Day | **102.3±7.24** | **147.2±8.58 ## **** | **151.6±6.68** |
| 6 | **Cholesterol (mg/dL)** | 0 Day | 46.8±6.74 | 45.9±6.85 | 17.3±3.17 |
| | | 30 Day | 48.1±6.49 | 40.3±6.76 | 17.8±4.21 |
| | | 60 Day | 49.1±49.1 | 37.6±5.88 | 18.4±4.33 |
| | | 90 Day | 50.4±50.4 | 34.9±7.13 | 19.2±4.12 |
| | | 125 Day | 53.7±53.7 | 32.0±6.57 | 21.0±3.78 |
| | | 155 Day | **56.6±56.6** | **28.1±5.45 ### **** | **23.0±3.75** |

| | | | | | |
|---|---|---|---|---|---|
| **7** | **Creatinine (mg/dL)** | 0 Day | 1.08±0.10 | 1.07±0.12 | 0.29±0.02 |
| | | 30 Day | 1.06±0.09 | 1.02±0.13 | 0.35±0.04 |
| | | 60 Day | 1.31±0.24 | 0.87±0.11 | 0.39±0.03 |
| | | 90 Day | 1.54±0.26 | 0.80±0.10 | 0.44±0.04 |
| | | 125 Day | 1.78±0.22 | 0.70±0.08 | 0.50±0.03 |
| | | 155 Day | **2.03±0.33** | **0.63±0.08 ### *** | **0.54±0.02** |
| **8** | **BUN (mg/dL)** | 0 Day | 16.23±1.21 | 16.19±0.92 | 4.03±0.13 |
| | | 30 Day | 16.36±1.17 | 15.26±0.74 | 4.07±0.15 |
| | | 60 Day | 16.66±1.14 | 14.57±0.66 | 4.14±0.15 |
| | | 90 Day | 16.80±1.17 | 13.27±0.78 | 4.22±0.15 |
| | | 125 Day | 16.99±1.20 | 11.05±0.79 | 4.32±0.16 |
| | | 155 Day | **17.11±1.22** | **8.94±0.73 ### ***** | **4.46±0.15** |
| **9** | **SGPT (IU/L)** | 0 Day | 34.30±1.65 | 34.12±1.91 | 22.20±1.48 |
| | | 30 Day | 34.06±1.42 | 32.20±1.52 | 23.25±1.52 |
| | | 60 Day | 34.93±1.21 | 30.78±1.41 | 24.56±1.43 |
| | | 90 Day | 36.02±1.18 | 29.78±1.03 | 25.70±1.78 |
| | | 125 Day | 37.16±1.11 | 28.87±0.89 | 27.05±1.84 |
| | | 155 Day | **38.29±1.23** | **28.03±0.76 ### *** | **27.56±1.52** |
| **10** | **SGOT (IU/L)** | 0 Day | 86.24±3.77 | 86.79±2.35 | 41.53±1.93 |
| | | 30 Day | 90.02±3.95 | 80.74±2.15 | 44.86±1.87 |
| | | 60 Day | 92.41±3.69 | 75.26±2.28 | 45.39±1.78 |
| | | 90 Day | 94.97±3.87 | 66.39±3.17 | 47.55±2.41 |
| | | 125 Day | 96.18±3.33 | 60.60±1.22 | 50.39±2.36 |
| | | 155 Day | **97.45±2.38** | **54.13±1.85 ### **** | **53.54±2.43** |
| **11** | **Total protein (g/dl)** | 0 Day | 7.59±1.06 | 7.75±0.88 | 4.74±0.95 |
| | | 30 Day | 8.22±1.07 | 7.55±0.78 | 5.15±0.73 |
| | | 60 Day | 8.73±0.73 | 7.39±0.80 | 5.57±0.77 |
| | | 90 Day | 9.83±0.59 | 7.15±0.94 | 5.81±0.87 |
| | | 125 Day | 10.96±0.35 | 7.04±0.88 | 6.56±0.80 |
| | | 155 Day | **12.14±0.53** | **7.12±0.86 ##** | **7.01±0.86** |

**Supplementary Table 7:** Detailed vital organ biomarker measurements of rats at stated time points post-plasma fraction treatment. The measurements (with standard deviations) were from 6 rats per group. These average values formed the graphs in Figure 3.

| Row | Figure | Tissue | Epigenetic Clock | Old Control | Young Control | Old Treated | Perc.Rejuv. |
|---|---|---|---|---|---|---|---|
| 1 | Figure2A | Blood | Rat Pan | 1.852 | 0.851 | 0.907 | 51.0% |
| 2 | Figure2B | Liver | Rat Pan | 2.121 | 0.658 | 0.783 | 63.1% |
| 3 | Figure2C | Heart | Rat Pan | 1.568 | 0.852 | 0.946 | 39.7% |
| 4 | Figure2D | Hypothalamus | Rat Pan | 1.650 | 0.822 | 1.540 | 6.6% |
| 5 | Figure2E | Blood | Rat Blood | 2.054 | 0.973 | 0.912 | 55.6% |
| 6 | Figure2F | Liver | Rat Liver | 2.363 | 0.685 | 0.727 | 69.2% |
| 7 | Figure2G | Heart | Rat Blood | 0.743 | 0.368 | 0.392 | 47.3% |
| 8 | Figure2H | Hypothalamus | Rat Brain | 2.350 | 1.012 | 1.983 | 15.6% |
| 9 | Figure2I | Blood | HumanRat Pan Age | 1.384 | 0.698 | 0.613 | 55.7% |
| 10 | Figure2J | Liver | HumanRat Pan Age | 1.944 | 0.644 | 0.364 | 81.3% |
| 11 | Figure2K | Heart | HumanRat Pan Age | 1.598 | 0.414 | 0.416 | 74.0% |
| 12 | Figure2L | Hypothalamus | HumanRat Pan Age | 1.684 | 0.562 | 1.666 | 1.1% |
| 13 | Figure2M | Blood | RelativeAge HumanRat | 0.316 | 0.184 | 0.169 | 46.7% |
| 14 | Figure2N | Liver | RelativeAge HumanRat | 0.351 | 0.133 | 0.070 | 79.9% |
| 15 | Figure2O | Heart | RelativeAge HumanRat | 0.438 | 0.185 | 0.228 | 48.0% |
| 16 | Figure2P | Hypothalamus | RelativeAge HumanRat | 0.428 | 0.164 | 0.343 | 19.9% |
| 17 | SuppFig12A | Blood | Rat Pan | 1.987 | 0.710 | 0.811 | 59.2% |
| 18 | SuppFig12B | Liver | Rat Pan | 2.183 | 0.530 | 0.642 | 70.6% |
| 19 | SuppFig12C | Heart | Rat Pan | 1.875 | 0.700 | 0.869 | 53.7% |
| 20 | SuppFig12D | Hypothalamus | Rat Pan | 1.937 | 0.707 | 1.509 | 22.1% |
| 21 | SuppFig12E | Blood | Rat Blood | 1.944 | 0.611 | 0.741 | 61.9% |
| 22 | SuppFig12F | Liver | Rat Liver | 2.041 | 0.612 | 0.628 | 69.2% |
| 23 | SuppFig12G | Heart | Rat Blood | 0.758 | 0.414 | 0.426 | 43.8% |
| 24 | SuppFig12H | Hypothalamus | Rat Brain | 2.056 | 0.659 | 1.620 | 21.2% |
| 25 | SuppFig12I | Blood | HumanRat Pan Age | 1.992 | 0.685 | 0.590 | 70.4% |
| 26 | SuppFig12J | Liver | HumanRat Pan Age | 2.175 | 0.597 | 0.403 | 81.5% |
| 27 | SuppFig12K | Heart | HumanRat Pan Age | 2.024 | 0.574 | 0.606 | 70.1% |
| 28 | SuppFig12L | Hypothalamus | HumanRat Pan Age | 1.929 | 0.621 | 1.707 | 11.5% |
| 29 | SuppFig12M | Blood | RelativeAge HumanRat | 0.533 | 0.166 | 0.157 | 70.6% |
| 30 | SuppFig12N | Liver | RelativeAge HumanRat | 0.552 | 0.153 | 0.114 | 79.4% |
| 31 | SuppFig12O | Heart | RelativeAge HumanRat | 0.546 | 0.155 | 0.210 | 61.6% |
| 32 | SuppFig12P | Hypothalamus | RelativeAge HumanRat | 0.525 | 0.169 | 0.415 | 20.9% |

**Supplementary Table 8: Mean values of the DNAm age estimates of the six epigenetic clocks in the plasma treatment study.** The columns report the figure panels of the corresponding barplot. The tissue that was used to evaluate the clock. The type of clock. The percentage of rejuvenation (last column) was calculated as follows: 100*(1-Old Treated/OldControl). The results for Figure 2 involved the preliminary versions of the six rat clocks. The results for Supplementary Figure 12 involved the final versions of the six rat clocks as detailed in Methods.

According to the six epigenetic clocks, the plasma fraction treatment rejuvenated liver by 73.4% (ranging from 63% to 81% depending on the clock, Supplementary Table 8), blood by 52% (ranging from 47 to 56%), heart by 52% (ranging from 40 to 74%), and hypothalamus by 11% (ranging from 1 to 20%). The rejuvenation effects are even more pronounced if we use the final versions of our epigenetic clocks: liver 75%, blood 66%, heart 57%, hypothalamus 19%. According to the final version of the eigenetic clocks, the average rejuvenation across four tissues was 54.2%.
